# Structural basis of broad protection against influenza virus by a human antibody targeting the neuraminidase active site via a recurring motif in CDR H3

**DOI:** 10.1101/2024.11.26.625467

**Authors:** Gyunghee Jo, Seiya Yamayoshi, Krystal M. Ma, Olivia Swanson, Jonathan L. Torres, James A. Ferguson, Monica L. Fernández-Quintero, Jiachen Huang, Jeffrey Copps, Alesandra J. Rodriguez, Jon M. Steichen, Yoshihiro Kawaoka, Julianna Han, Andrew B. Ward

## Abstract

Influenza viruses evolve rapidly, driving seasonal epidemics and posing global pandemic threats. While neuraminidase (NA) has emerged as a vaccine target, shared molecular features of NA antibody responses are still not well understood. Here, we describe cryo-electron microscopy structures of the broadly protective human antibody DA03E17, which was previously identified from an H1N1-infected donor, in complex with NA from A/H1N1, A/H3N2, and B/Victoria-lineage viruses. DA03E17 targets the highly conserved NA active site using its long CDR H3, which features a DR (Asp–Arg) motif that engages catalytic residues and mimics sialic acid interactions. We further demonstrate that this motif is conserved among several NA active site-targeting antibodies, indicating a common receptor mimicry strategy. We also identified potential antibody precursors containing this DR motif in all donors of a healthy human donor BCR database, highlighting the prevalence of this motif and its potential as vaccine targeting. Our findings reveal shared molecular features in NA active site-targeting antibodies, offering insights for NA-based universal influenza vaccine design.

## Introduction

Influenza viruses remain a significant global health threat, causing an estimated 290,000 to 650,000 seasonal influenza-associated respiratory deaths and 3 to 5 million cases of severe illness each year^1^. Despite a temporary decline during the COVID-19 pandemic^2^, influenza cases and fatalities have returned to pre-pandemic levels^3,4^. Currently, both influenza A viruses (IAVs, H1N1 and H3N2) and influenza B viruses (IBVs, Victoria-lineage) are co-circulating globally, while the B/Yamagata lineage has not been reliably detected since March 2020^5,6^. Influenza A viruses, in particular, are known for their rapid antigenic evolution, leading to the emergence of new variants that may impact antibody recognition and pose new pandemic threats^7,8^. Recent antigenic drift in A/H3N2 viruses has introduced an N-glycosylation site at residue 245 on the surface glycoprotein neuraminidase (NA) since the 2014/2015 season, which can shield certain epitopes and is known to reduce the activity of monoclonal and human serum NA-specific antibodies^9–11^. Additionally, a highly pathogenic avian influenza (HPAI) H5N1 strain is currently spreading among cattle and other mammals in the United States^12,13^. This unprecedented outbreak has resulted in virus spillover into humans, raising concerns about zoonotic transmission and potential pandemic risks^14,15^.

NA is one of the two surface glycoproteins of influenza viruses, responsible for cleaving sialic acid from glycoproteins on the host cell surface, facilitating the release of progeny viruses^16^, while hemagglutinin (HA) mediates viral entry by binding to these sialic acid receptors^17^. Due to its essential role in the viral life cycle, the catalytic activity of NA has been the target of small-molecule drugs such as oseltamivir (Tamiflu)^18^. However, current influenza vaccines primarily induce antibody responses against HA, which can reduce disease severity but often fail to prevent infection due to rapid antigenic drift in HA, necessitating annual updates^19,20^. In contrast, antigenic drift in NA occurs at a slower rate and independently from HA^21,22^. Antibodies targeting NA are an independent correlate of protection with broader cross-reactivity^23,24^, making NA a promising vaccine target for inducing protective antibodies less sensitive to seasonal drift.

Identifying recurring molecular features of broadly protective antibodies across different individuals is crucial for designing effective immunogens in the development of universal influenza vaccines^25,26^. The highly conserved HA stem is frequently targeted by multidonor class antibodies with shared sequence features across individuals^27–31^. Structural insights from these antibodies have been instrumental in advancing universal influenza vaccine development^32,33^. In recent years, several broadly protective antibodies against NA have been identified^34–39^. Notably, antibodies targeting the highly conserved enzymatic active site of NA exhibit broader cross-reactivity against diverse influenza strains compared to other NA-specific antibodies and have provided crucial insights for the development of NA-based universal influenza vaccines^40–44^. However, our understanding of broadly protective NA epitopes remains limited compared to HA^45^, emphasizing the need to identify more broadly protective antibodies. This would allow shared molecular features of NA antibody responses to converge into clearer patterns, guiding the development of effective universal vaccines.

We previously identified and functionally characterized the human monoclonal antibody (mAb) DA03E17, which was isolated from an individual infected with the A/H1N1pdm09 virus during the 2015–2016 influenza season^43^. DA03E17 exhibited broad cross-reactivity against NAs from both group 1 and group 2 IAVs, as well as both lineages of IBVs. DA03E17 bound to or near the enzymatic active site of NA, effectively inhibiting sialidase activity, neutralizing diverse influenza strains *in vitro*, and providing protection *in vivo* across multiple subtypes of IAVs and IBVs. However, the precise epitope targeted by DA03E17 was not defined. In this study, we present the cryo-electron microscopy (cryo-EM) structures of DA03E17 in complex with NA from A/H1N1, A/H3N2, and B/Victoria-lineage viruses. DA03E17 retained binding to NAs with oseltamivir-resistant mutations and showed strong affinity for NAs from recent H3N2 strains and the Bovine HPAI H5N1 virus, demonstrating its potential to target both seasonal and emerging zoonotic viruses. Structural analysis revealed that DA03E17 engages the highly conserved NA active site using its long complementarity-determining region (CDR) H3, where the DR motif mimics the sialic acid receptor by interacting with key catalytic residues. This receptor mimicry strategy, shared by several NA active site-targeting antibodies, suggests a convergent mechanism for broad reactivity across influenza A and B viruses, offering insights for the development of NA-based universal influenza vaccines.

## Results

### Binding properties of DA03E17 to antiviral-resistant variants and diverse Influenza NAs

We previously isolated and characterized a human mAb DA03E17 from peripheral blood mononuclear cells (PBMCs) of an individual who was infected with A/H1N1pdm09 virus in the 2015–2016 influenza season^43^. DA03E17 showed broad cross-reactivity against NAs from IAV group 1 (N1, N4, N5, N8), group 2 (N2, N3, N6, N7, N9), and IBVs (**Extended Data Fig. 1a**)^43^. Furthermore, DA03E17 inhibited the sialidase activity of NA, neutralized both IAVs and IBVs *in vitro*, and provided *in vivo* protection against several subtypes of influenza virus^43^. DA03E17 uses the IGHV4-31 and IGKV1-12 heavy and light chain V genes, respectively, and has a long CDR H3 (19 amino acids in the IMGT CDR definition scheme) (**Extended Data Fig. 2**). In addition to the previously reported broad reactivity of DA03E17, here we further characterized its ability to retain binding to NAs containing oseltamivir-resistant mutations. We generated recombinant N1 NA from H1N1 A/Brisbane/02/2018 (BB18), which contains previously reported stabilizing mutations derived from a computationally-designed NA (stabilized NA protein, sNAp) in the inter-protomeric interface to maintain a closed tetrameric state^46^. We produced BB18 N1 sNAp containing either the major oseltamivir-resistance mutation H274Y^47^ or other resistance substitutions, I222V or S246N (N2 numbering)^48^. Additionally, we generated recombinant N2 NA from H3N2 A/Indiana/10/2011 (IN11) with either the E119V or I222L substitution^49^. We measured the binding of DA03E17 to these recombinant N1 and N2 NAs by ELISA. While DA03E17 retained its binding to BB18 N1 sNAp with either the H274Y, I222V, or S246N substitution, it exhibited reduced binding to IN11 N2 with either the E119V or I222L substitution, although some level of binding was still observed (**Extended Data Fig. 1b**). Biolayer interferometry (BLI) indicated that DA03E17 bound to recombinant NAs from H1N1 A/California/07/2009 (CA09 N1 sNAp), H3N2 A/Perth/16/2009 (PT09 N2), and H3N2 A/Indiana/08/2011 (IN11 N2) with sub-picomolar affinity, while it bound to NAs from B/Colorado/06/2017 (B/Victoria-lineage; CO17 B) and more recent H3N2 strains, including A/Kansas/14/2017 (KS17 N2) and A/Hong Kong/2671/2019 (HK19 N2), with sub-nanomolar to weaker nanomolar affinities (**Extended Data Fig. 3**). Notably, DA03E17 also bound to recombinant N1 sNAp derived from the HPAI H5N1 clade 2.3.4.4b virus with nanomolar affinity, which is currently spreading across dairy herds and other mammals in 14 states in the United States^12–14^ (**Extended Data Fig. 3**).

### Cryo-EM structures of DA03E17 Fab in complex with N1, N2, and B NAs

To elucidate the epitope of DA03E17 and the structural basis for its broad cross-reactivity, we determined the cryo-EM structures of the DA03E17 Fab in complex with CA09 N1 sNAp, and KS17 N2 and CO17 B NAs at 2.67, 2.86, and 2.47 Å resolution, respectively (**Extended Data Fig. 4** **and Table 1**). Additionally, a cryo-EM structure of KS17 N2 NA was determined in its apo-form at 2.75 Å resolution using the same data set of DA03E17-KS17 N2 NA complex (**Extended Data Fig. 4**). In all three complex structures, DA03E17 binds in a similar orientation to all three NAs, with each Fab interacting with just one protomer of the NA tetramer (**Fig. 1a–c and Extended Data Fig. 5a**–f). DA03E17 fully blocks the NA active site by protruding the CDR H3 into the active site pocket (**Fig. 1c and Extended Data Fig. 5c**,f), consistent with our previous study using the small molecule-based NA-Star assay, which demonstrated that the NI activity of DA03E17 is due to direct inhibition^43^. No large conformational changes in the global structure of the NA protein are observed compared with corresponding wild-type structures, as indicated by an RMSD of 0.356, 0.402, and 0.314 Å across all pairs for the N1, N2, and B NAs, respectively (**Extended Data Fig. 5g**–i).

**Fig. 1:**
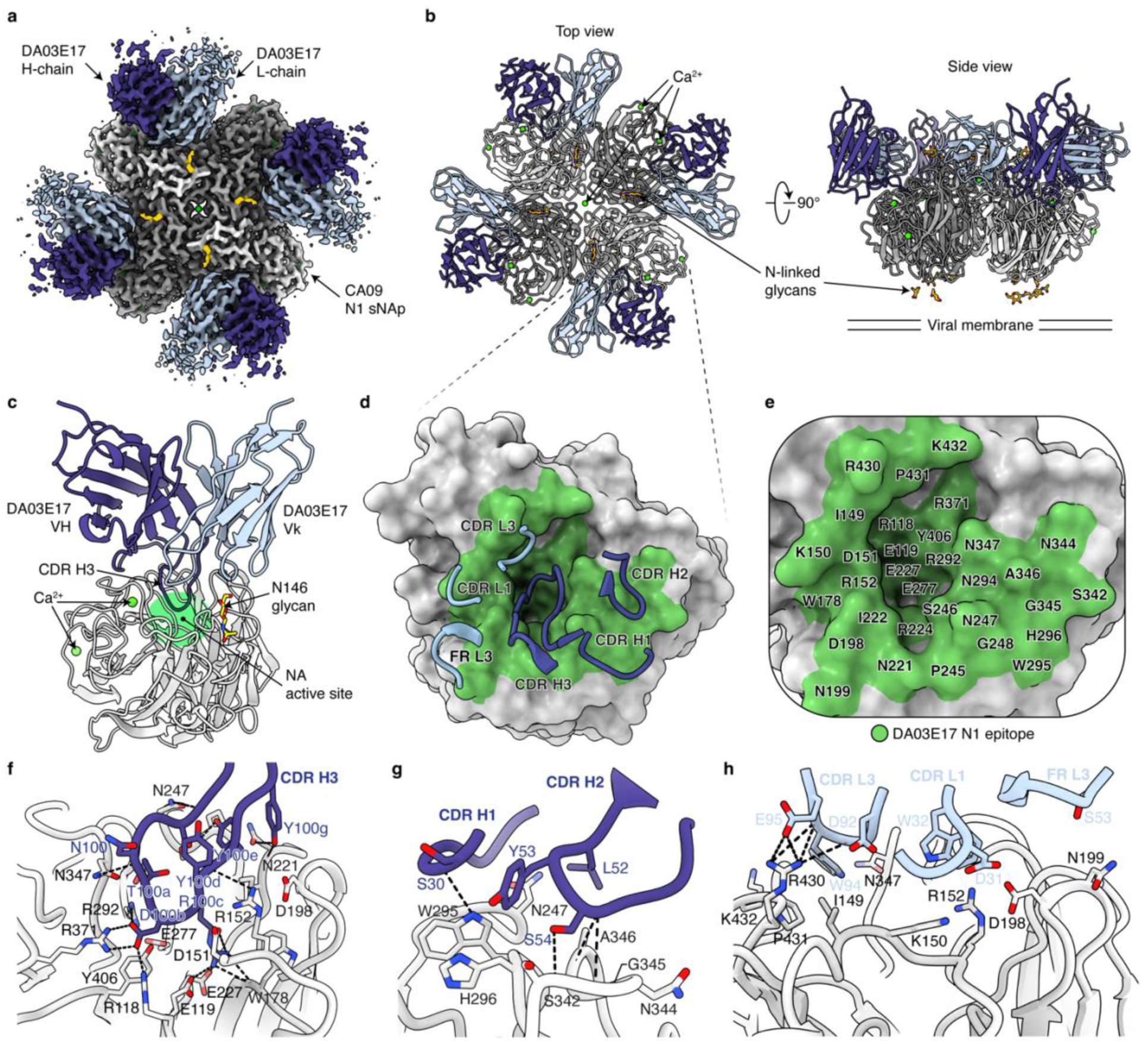
Cryo-EM structure of DA03E17 Fab in complex with CA09 N1 sNAp. **a**,**b**, Cryo-EM map at 2.67 Å resolution (**a**) and two orthogonal views of the atomic model from the top and side (**b**); only the Fab variable region is built into the density. DA03E17 heavy chain, dark blue; light chain, light blue; NA head domain tetramer; shades of gray; NA glycans, gold; Ca ^2+^ ion, lime. **c**, Ribbon diagram of the CA09 N1 sNAp protomer bound to one DA03E17 Fab. The NA active site targeted by CDR H3 is highlighted by a green oval. VH, heavy chain variable domain; Vk, kappa light chain variable domain. **d**,**e**, DA03E17 epitope mapped on the NA protomer surface with CDR loops involved (**d**) or residue labels (**e**). The DA03E17 N1 epitope is highlighted in green. **f**–**h**, Detailed interactions between CA09 N1 sNAp and DA03E17 CDR H3 (**f**), CDR H1 and H2 (**g**), and CDR L1, L3, and FR L3 (**h**).

**Table 1.**
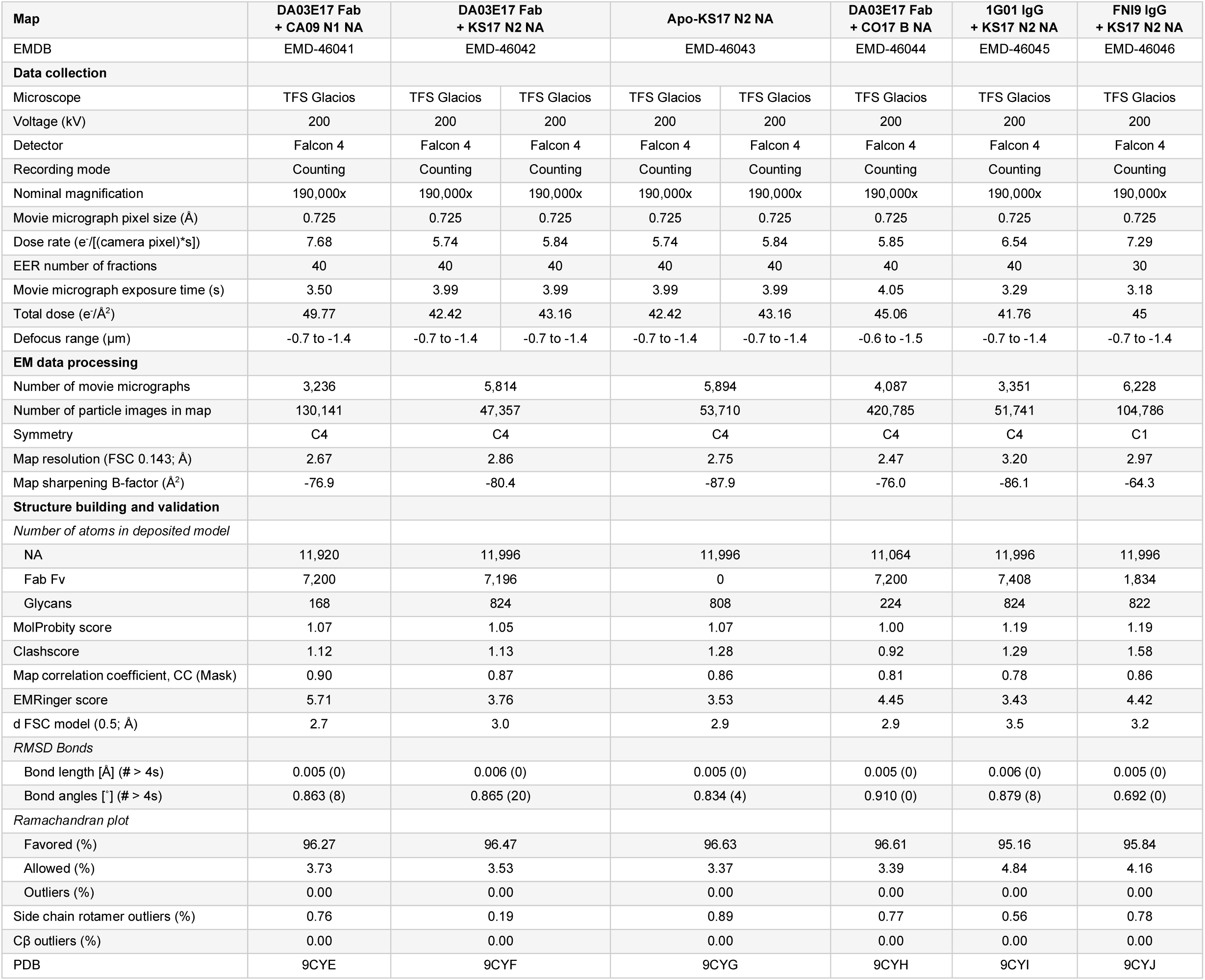
Cryo-EM data collection, processing, model refinement and validation statistics.

In the DA03E17-CA09 N1 sNAp complex structure, the buried surface area (BSA) of the DA03E17 and N1 NA interface is 1022.2 Å^2^, with the heavy chain accounting for 78% of the interaction. DA03E17 interacts with CA09 N1 sNAp using all CDR loops except CDR L2 (**Fig. 1d**), and the DA03E17 epitope on CA09 N1 sNAp consists of 32 residues around the active site pocket (**Fig. 1e**). The 19-residue CDR H3 contributes most of the Fab interactions with 19 N1 NA residues including seven catalytic site residues (R118, D151, R152, R224, R292, R371, and Y406) and six framework residues (E119, W178, I222, E227, E277, and N294) (**Fig. 1f**) which are strictly conserved across influenza A group 1, group 2, and influenza B NAs (**Extended Data Table 1**). The CDRs H1, H2, L1, L3, and framework region (FR) L3 predominantly interact with residues located at the periphery of the active site (I149, K150, D198, N199, N247, W295, H296, S342, N344, G345, A346, N347, R430, P431, and K432), further strengthening binding (**Fig. 1g,h**). Overall, DA03E17 utilizes both heavy and light chain CDRs to bind the active site of NA, and a total of 24 hydrogen bonds were observed between DA03E17 and CA09 N1 sNAp. In the DA03E17-KS17 N2 and DA03E17-CO17 B NA complex structures, the BSAs at the interfaces are 1224.5 Å² and 977.6 Å², respectively, indicating a slightly larger interaction surface with the KS17 N2 NA compared to the CA09 N1 and CO17 B NAs. Most of the interactions mediated by the CDR H3 in the DA03E17-KS17 N2 and DA03E17-CO17 B NA complexes are similar to those observed in the CA09 N1 sNAp complex (**Extended Data Fig. 6a**,e), with key epitope residues being conserved across IAV and IBV NAs (**Extended Data Table 1**), maintaining a consistent binding mode. These data establish that DA03E17 directly targets the highly conserved residues in the enzymatic active site of NA using its long CDR H3, thereby blocking the active site and inhibiting sialidase activity of NA.

### DA03E17 targets highly conserved epitopes in N1, N2, and B NAs

To further understand the broad cross-reactivity of DA03E17, we examined the sequence conservation of DA03E17 epitopes on NAs from H1N1, H3N2, and IBV viruses that have been circulating for several decades. (**Fig. 2a–c**). The epitopes of DA03E17 are well conserved among human seasonal H1N1 (circulating from 1977 to 2023) and H3N2 (from 1968 to 2023) IAVs, as well as Victoria/2/87-like IBVs (from 1987 to 2023). The majority of epitope residues are highly conserved or conservatively substituted on H1N1 (25 out of 32, 78%), H3N2 (25 out of 35, 71%), and B/Vic (25 out of 30, 83%) NAs (**Fig. 2d**). Substitutions at positions 199 and 221—the two most variable residues within the N1 and N2 epitopes—from N199 (CA09) to S199 (H1N1 A/Yokohama/94/2014) and N221 (CA09) to K221 (H1N1 A/Puerto Rico/8/1934) in N1, as well as from K199/D221 (KS17) to E199/K221 (H3N2 A/Fujian/411/2002; FJ02) in N2, did not compromise the NI activity of DA03E17, as shown in our previous study^43^. Similarly, other substitutions in the N2 epitope, from N147/R150/K344 (KS17) to D147/H150/E344 (FJ02), also did not impair the NI activity of DA03E17^43^, suggesting that DA03E17 binding is resilient to these substitutions. The core of DA03E17 epitope in the active site, which is targeted by the CDR H3, is strictly conserved across A/H1N1, A/H3N2, and B/Victoria-like viruses due to the functional constraints related to sialic acid receptor recognition (**Fig. 2d,e**). Therefore, the broad cross-reactivity of DA03E17 can be partially explained by the strict conservation or conservative substitution of key epitope residues, along with its resilience to substitutions around the NA active site. Notably, several conserved residues in the active site pocket are charged amino acids, forming distinct charged patches with R118, R292, and R371 contributing to positive charges, and E119, D151, E227, and E277 forming negative charges (**Fig. 2e,f**). D100b and R100c, which form the DR (Asp–Arg) motif at the tip of the DA03E17 CDR H3, have an electrostatic surface that is complementary to these charged patches, enabling extensive salt bridge formation for targeted interaction (**Figs. 2g, 1f, and Extended Data Fig. 6a**,e). Overall, we found that the DA03E17 epitope is largely conserved, especially the active site residues targeted by the CDR H3, which are strictly conserved across the NAs of H1N1, H3N2, and B/Victoria-like viruses that have been circulating for several decades, highlighting the critical role of the CDR H3 in the broad cross-reactivity of DA03E17.

**Fig. 2:**
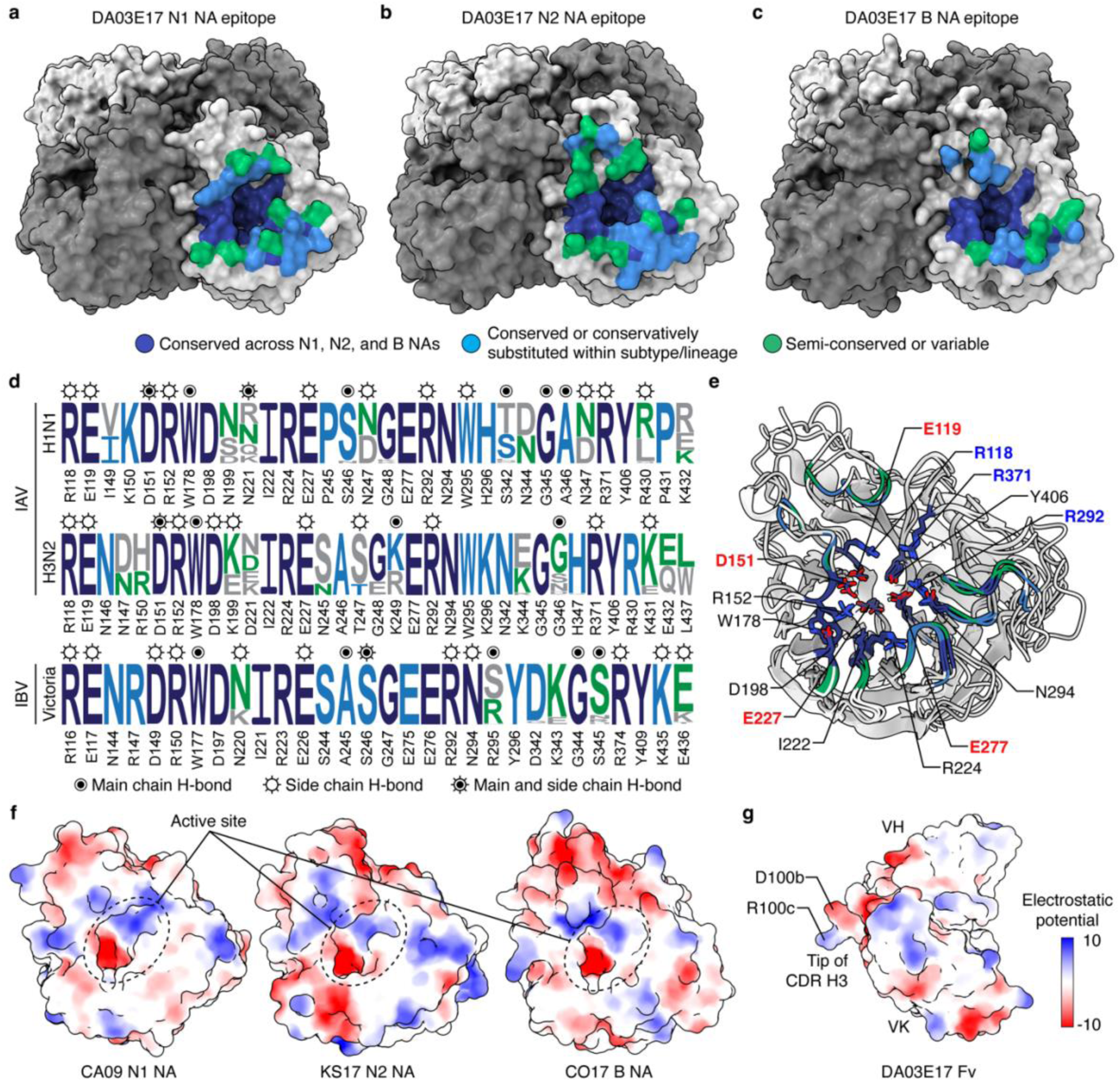
Sequence conservation of the DA03E17 epitope. **a**–**c**, Sequence conservation of DA03E17 epitope mapped onto surface representations of CA09 N1 (**a**), KS17 N2 (**b**), and CO17 B (**c**) NAs. NA residues conserved across A/H1N1, A/H3N2, and B/Victoria-like viruses are colored in dark blue, while those conserved or conservatively substituted within subtype and lineage are colored in sky blue. Semi-conserved or variable residues are colored in green. **d**, Conservation of DA03E17 epitopes based on NA sequences from human seasonal H1N1 (circulating from 1977 to 2023) and H3N2 (1968–2023) IAVs and Victoria/2/87-like IBVs (1987– 2023). Circled bullets denote NA main chain contacts; open circles with rays denote side chain contacts; Circled bullets with rays denote both main and side chains contacts. **e**, Superimposed protomers of CA09 N1 sNAp, and KS17 N2 and CO17 B NAs with conserved active site residues shown as sticks and critical charged residues highlighted. **f**,**g**, Surface representation of CA09 N1, KS17 N2, and CO17 B NA protomers (**f**) and DA03E17 Fv (**g**) colored by electrostatic potential from −10 to +10 *k*_B_*T*/*e*_c_, with dotted circles indicating charged patches in the NA active site pocket.

### DA03E17 accommodates the N-glycans of the recently circulating human H3N2 viruses

The NA of circulating human A/H3N2 viruses has undergone substantial antigenic drift since 2014, resulting from mutations at positions 245 and 247 that introduced an N-linked glycan near the active site^9^ (**Fig. 3a and Extended Data Fig. 7a**). By the 2016/17 season, nearly all circulating H3N2 viruses carried this glycan, which has become fixed in recent strains and has been shown to shield the NA active site, reducing the efficacy of certain NA active site-targeting antibodies ^9–11,44^. Our previous study showed that DA03E17 maintained inhibitory activity against the NAs of recently circulating H3N2 viruses, despite the presence of the N245 glycan^43^. BLI analysis revealed that while DA03E17 exhibited reduced binding affinity for drifted N2 NAs (KS17 and HK19 N2) carrying the N245 glycan, as well as mutant IN11 N2 NA (IN11 N2 NAT) with an introduced N245 glycosylation site, compared to NAs without the glycan, it still maintained nanomolar affinities (**Fig. 3b and Extended Data Fig. 3**). These results indicate that the N245 glycan may affect the binding of DA03E17 to the NA active site. To investigate in detail how the N245 glycan affects the binding of DA03E17, we compared the structure of the DA03E17-KS17 N2 NA complex with that of the apo-KS17 N2 NA, which was determined from the same data set of the DA03E17-KS17 N2 NA complex (**Extended Data Fig. 4**). In the apo-KS17 N2 structure, the N245 glycan would cause a steric clash with DA03E17 CDR H1, while the N146 glycan would clash with CDR L1. However, upon binding to KS17 N2 NA, DA03E17 induces conformational changes in both the N245 and N146 glycans, allowing it to maintain binding by repositioning these glycans and avoiding the clashes (**Fig. 3c and Extended Data Fig. 7b**).

**Fig. 3:**
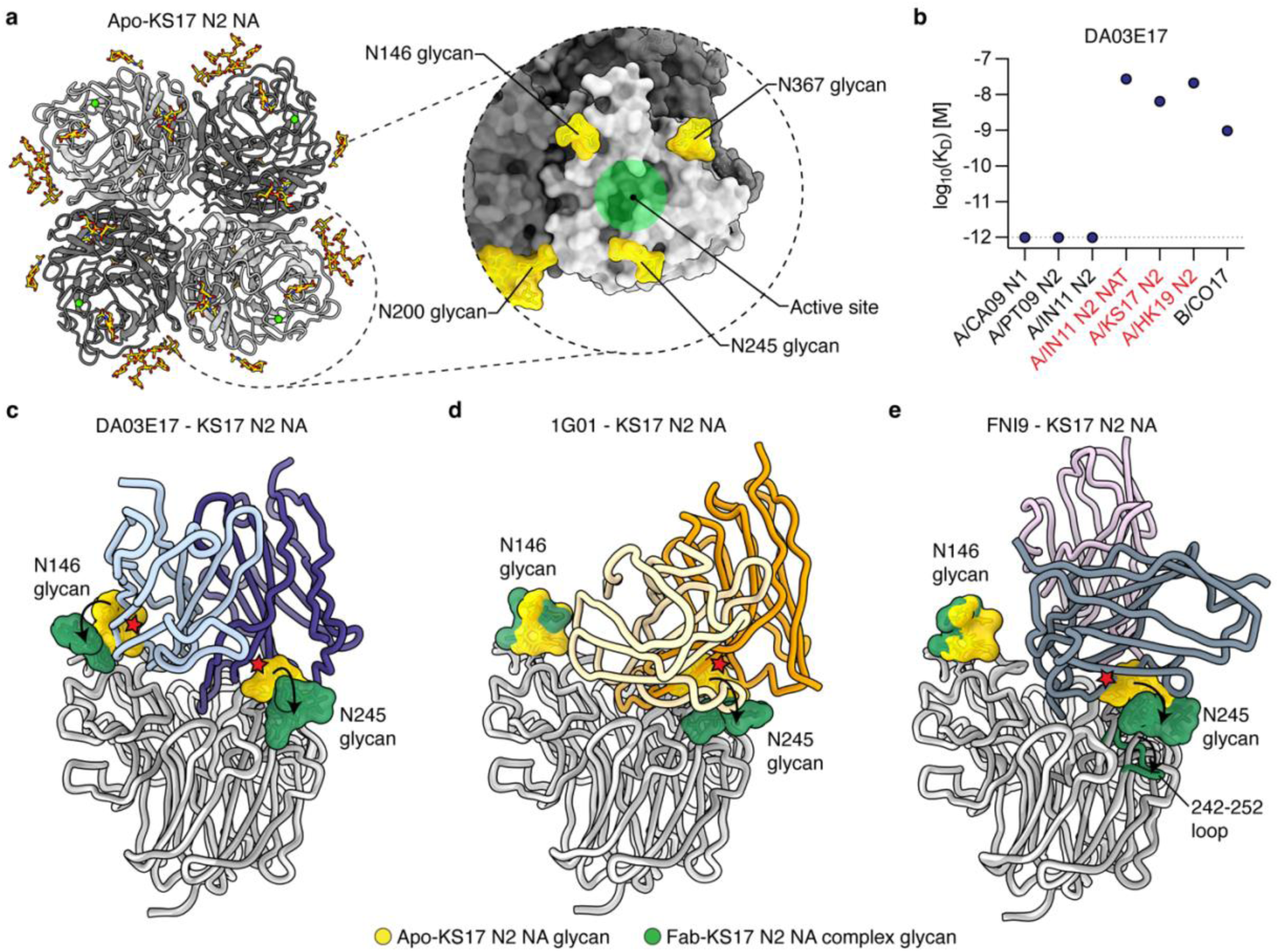
DA03E17 accommodates the N-glycans of drifted N2 NA. **a**, Cryo-EM structure of KS17 N2 NA in apo-form at 2.75 Å resolution. The inset shows a magnified view of a KS17 N2 NA protomer, with the active site indicated by a green circle, and the surrounding glycans are shown in yellow. **b**, Binding affinity of DA03E17 IgG to recombinant NAs, measured by BLI with CA09 N1, PT09 N2, IN11 N2, IN11 N2 S245N/S247T (NAT) mutant, KS17 N2, HK19 N2, and CO17 B NAs. NAs with an N-glycan at position 245 are highlighted in red. K_D_ was estimated using a 1:1 binding model. Dotted line represents detection limit. **c**–**e**, Overlay of the apo-KS17 N2 NA protomer (gray) with Fab-bound KS17 N2 NA protomers (white): DA03E17 Fab (**c**), 1G01 Fab (**d**), and FNI9 Fab (**e**). The N146 and N245 glycans on the apo-KS17 N2 NA are shown in yellow, and those on the Fab-bound KS17 N2 NA are shown in green. Steric clash shown with red stars. Conformational changes in glycans and the 242–252 loop are indicated by arrows.

To further investigate the structural changes in the N245 and N146 glycans upon DA03E17 binding to KS17 N2 NA, we performed molecular dynamics (MD) simulations. In the apo-KS17 N2 NA structure, the MD simulations revealed that the N245 and N146 glycans formed clusters that obstructed access to the NA active site and potentially blocked the binding of DA03E17 (**Extended Data Fig. 7c**,d). In contrast, MD simulations of the DA03E17-KS17 N2 NA complex demonstrated that DA03E17 binding induced significant structural changes in both the N245 and N146 glycans. The glycans shifted to adopt specific conformations, moving away from the DA03E17 binding interface. These movements aligned with the cryo-EM structural data (**Fig. 3a,c**), further supporting that DA03E17 induces a rearrangement of the N245 and N146 glycans to facilitate stable binding to the NA active site. Such conformational changes in the glycans may require additional energy, potentially contributing to the reduced affinity of DA03E17 for N2 NAs carrying the N245 glycan. Considering that the N146 glycan is also present in other N1 and N2 NAs, such as CA09 N1, PT09 N2, and IN11 N2, where DA03E17 binds with sub-picomolar affinity (**Extended Data Fig. 3**), it is likely that the N245 glycan plays a more prominent role in the observed reduction in affinity.

Previous studies have also explored how NA active site-targeting antibodies maintain binding and protective efficacy against contemporary H3N2 viruses carrying the N245 glycan, offering insights into the mechanisms these antibodies employ to accommodate this glycan^11,44^. The broadly cross-reactive antibody, 1G01, was shown to maintain protective efficacy *in vivo* despite reduced inhibition against viruses harboring the N245 glycan. Although structural analysis using negative-stain electron microscopy indicated that 1G01 still binds the active site of N2 NA with the N245 glycan, the precise mechanism remained unclear. Another broadly cross-reactive antibody, FNI9, was shown to not only induce conformational changes in the N245 glycan but also alter the conformation of the 242–252 loop of HK19 N2 NA, which contains the N_245_AT_247_ glycosylation motif. To compare the N245 glycan accommodation mechanisms of 1G01 and FNI9 with that of DA03E17 directly, we determined the cryo-EM structures of the 1G01-KS17 N2 NA and FNI9-KS17 N2 NA complexes at 3.20 Å and 2.97 Å resolutions, respectively (**Extended Data Fig. 8**). Structural comparison revealed that CDR H3 of 1G01 would clash with the N245 glycan in the apo-KS17 N2 NA structure. However, similar to DA03E17, 1G01 induces conformational changes in the N245 glycan, shifting it away from the binding interface and maintaining binding to the NA (**Fig. 3d**). For FNI9, similar to previous findings with HK19 N2 NA^44^, structural comparison revealed that it induces conformational changes not only in the N245 glycan but also in the 242–252 loop of KS17 N2 NA (**Fig. 3e**). This rearrangement allows FNI9 to maintain binding, suggesting that its glycan accommodation mechanism is not HK19 N2 NA-specific but rather a broader adaptation to drifted N2 NAs carrying the N245 glycan in general. To summarize, the three broadly cross-reactive antibodies, DA03E17, 1G01, and FNI9, accommodate the N245 glycan on drifted N2 NA through two primary mechanisms. DA03E17 and 1G01 induce conformational changes in the glycan, shifting it away from the binding interface, while FNI9 not only induces glycan rearrangement but also alters the conformation of the 242–252 loop, indicating a more extensive structural rearrangement. These results demonstrate two distinct strategies employed by NA active site-targeting antibodies to maintain effective active site binding.

### DA03E17 mimics the interaction of sialic acid using a conserved motif in the CDR H3

The DA03E17 footprint on the NA spans the entire active site pocket, fully covering it and blocking access to critical residues involved in enzymatic activity (**Fig. 1e**), suggesting that DA03E17 may inhibit NA activity through a mechanism similar to known NA inhibitors, which block the active site by mimicking sialic acid interactions^50,51^. We compared the interaction formed between the DA03E17 CDR H3 and the NA active site residues with that of the sialic acid receptor and oseltamivir. Remarkably, the carboxylate side chain of D100b in the DA03E17 CDR H3 forms the same salt bridge interaction network with NA residues R118, R292, and R371 as the carboxylate group of sialic acid and oseltamivir (**Fig. 4a–c**). The side chain of R100c also forms contacts with NA residues D151 and E227, recapitulating those of sialic acid and oseltamivir, further indicating receptor mimicry by DA03E17 CDR H3. Such receptor mimicry by antibody CDR H3 has been previously observed in other NA active site-targeting antibodies^36,41,44^. In our structural comparison, we found that the DR (Asp–Arg) motif in DA03E17 functions similarly to those in the previously reported pan-influenza NA mAb FNI9^44^ and the broadly cross-reactive anti-IBV NA mAb 1G05^41^, mimicking sialic acid interactions and blocking the NA active site (**Fig. 4d,e**). Notably, the DR motif is reversed in order in FNI9, appearing as R–D instead of D–R, as seen in DA03E17 CDR H3 (**Fig. 4i**), yet both motifs mimic sialic acid interactions from nearly identical positions in the active site. Additionally, we identified the same sialic acid-mimicking DR motif in the previously reported cross-group mAb Z2B3 (**Fig. 4f**), which binds to both N1 and N9 NAs^42^. These findings highlight the conserved role of DR motifs in mediating receptor mimicry across multiple NA active site-targeting antibodies, supporting the idea of sequence and structural convergence in antibody responses to NA. Similarly, sialic acid mimicry has been previously reported in the N9 NA-specific mAb NA-45^36^, where the negatively charged side chain of E (Glu), similar to D (Asp), is used to mimic the carboxylate group of sialic acid (**Fig. 4g**). This type of sialic acid mimicry, using D (Asp) and E (Glu) to mimic the carboxylate group of sialic acid, has also been observed in several HA receptor-binding site (RBS)-targeting antibodies^52–57^, highlighting the critical role of negatively charged amino acids in mimicking the carboxylate group of sialic acid within both the HA RBS and the NA active site. We also compared the structure of another pan-influenza NA mAb, 1G01, which has an R (Arg) at the tip of its CDR H3, but its interactions were distinct from those mediated by the DR motif (**Fig. 4h**). In summary, these NA active site-targeting antibodies possess relatively long CDR H3 loops (**Fig. 4i**), and while the Fabs have varying angles of approach and the CDR H3 loops engage the active site pocket in diverse ways (**Fig. 4j,k**), the DR or RD motifs remain structurally conserved among antibodies with receptor mimicry.

**Fig. 4:**
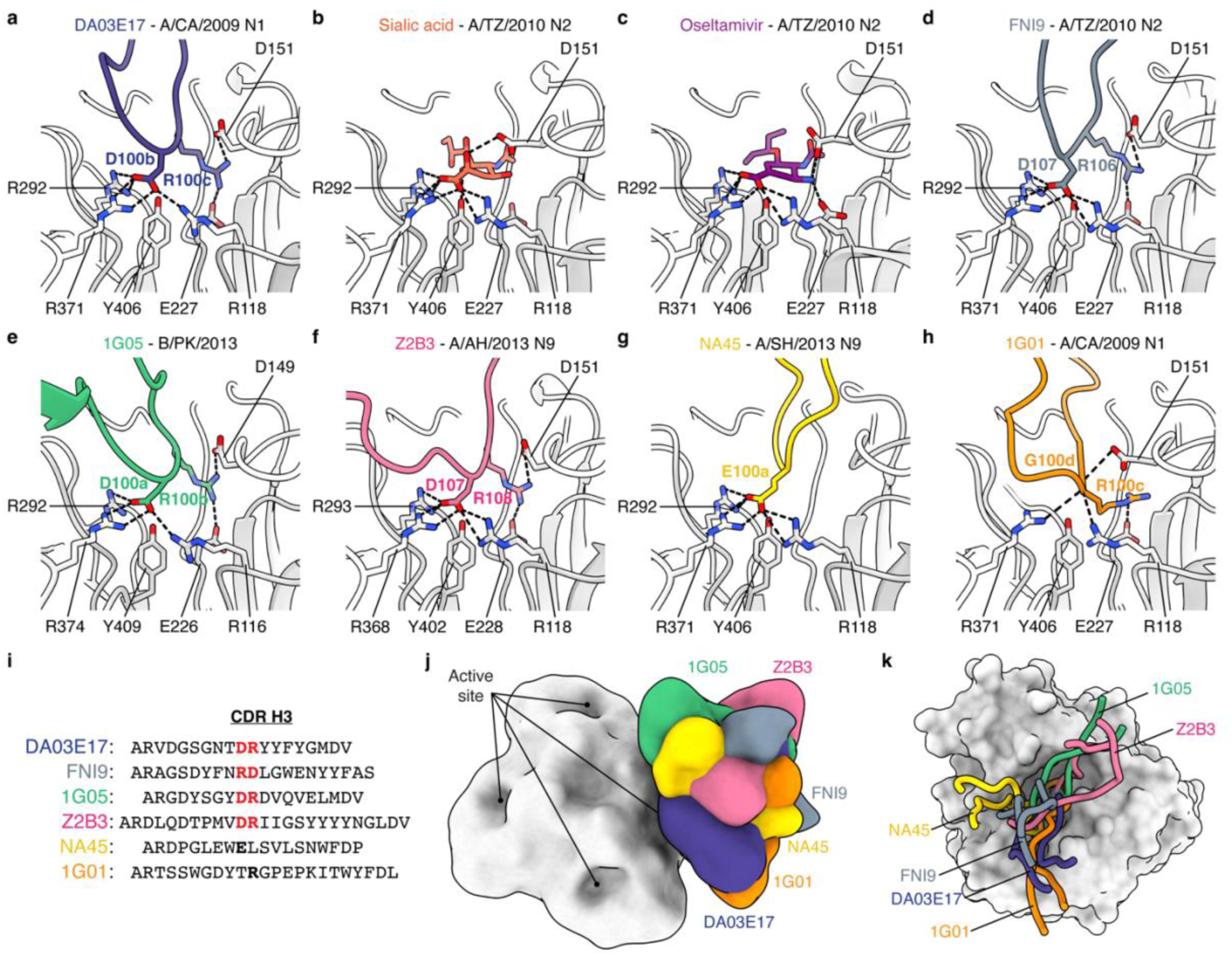
**The DR motif is a recurring molecular feature in the CDR H3 of NA active site**- **targeting antibodies. a**, Network of salt bridge interactions between D100b and R100c in the CDR H3 of DA03E17 and active site residues of CA09 N1 sNAp. **b**,**c**, Salt bridge interactions between sialic acid (**b**) and oseltamivir (**c**) with active site residues of H3N2 A/Tanzania/205/2010 (TZ10) NA (PDB: 4GZQ, 4GZP). **d**, Salt bridge interactions between D107 and R106 in the CDR H3 of FNI9 and TZ10 N2 NA (PDB: 8G3N). **e**, Salt bridge interactions between D100a and R100b in the CDR H3 of 1G05 and B/Phuket/3073/2013 NA (PDB: 6V4N). **f**, Salt bridge interactions between D107 and R108 in the CDR H3 of Z2B3 and H7N9 A/Anhui/1/2013 NA (PDB: 6LXJ). **g**, Salt bridge interactions between E100a in the CDR H3 of NA45 and H7N9 A/Shanghai/2/2013 NA (PDB: 6PZE). **h**, Hydrogen bonds and salt bridge interactions between R100c in the CDR H3 of 1G01 and CA09 N1 NA (PDB: 6Q23). **i**, Alignment of the CDR H3 sequences of DA03E17, FNI9, 1G05, Z2B3, NA45, and 1G01, with DR motifs mimicking the interaction of sialic acid highlighted in red. **j**,**k**, Overlay of low-pass filtered structures of NA active site-targeting antibodies bound to the tetrameric NA head domain (**j**), and a detailed view of their CDR H3 loops occupying the active site pocket of the NA protomer (**k**).

### DR motif precursors are prevalent in human antibody repertoire

To assess the feasibility of eliciting NA active site-targeting antibodies that mimic sialic acid binding through vaccination, we employed a bioinformatic approach to identify the prevalence of DR motif precursors in the human antibody repertoire. We utilized an ultradeep next-generation sequencing (NGS) dataset of 1.1 x 10^9^ antibody heavy chain sequences from 14 healthy human donors for the precursor frequency search^58,59^. Structural analysis of six NA active site-targeting antibodies indicated that a long CDR H3 loop was necessary to access the highly conserved residues within the active site pocket (**Fig. 4k**). A crucial feature of four cross-reactive antibodies was a centrally located DR or RD motif within the long CDR H3 loop that mimicked the interactions of the sialic acid receptor (**Figs. 4 and 5a**). Notably, in all three of the DR-motif antibodies for which genetic sequences were available (DA03E17 and 1G05) or the D gene position could be predicted (Z2B3), the DR motif was encoded at least partially by non-templated junction residues between the D and J genes (**Fig. 5b**), suggesting it could be a limiting factor for DR motif precursor frequency. Guided by these findings, we screened for antibodies with long CDR H3 loops (19–30 amino acids) containing a centrally located DR or RD motif, flanked by at least 8 amino acids on either side (**Fig. 5c**). Such DR or RD motifs BCRs were identified in all 14 donors, with frequencies of 1 in 1,143 and 1 in 742, respectively (**Fig. 5d**). These BCRs exhibited diversity in their D gene utilization, with no single D gene dominating the pool. While the proportion of these DR or RD motif BCRs capable of maturing into functional NA antibodies remains to be determined, the substantial prevalence of these precursors suggests they hold promise as vaccine targets.

**Fig. 5:**
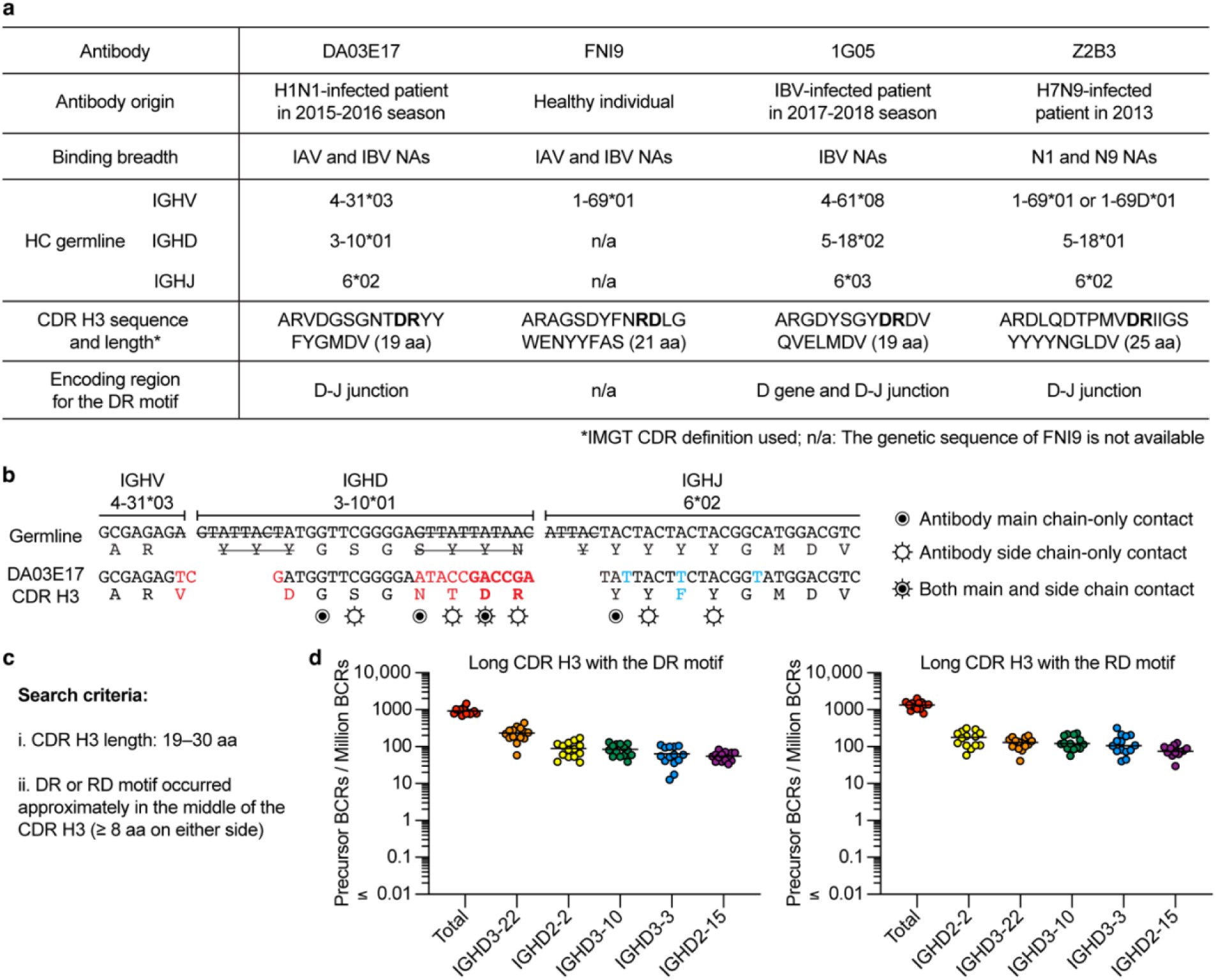
DR motif precursors are prevalent in humans. **a**, Genetic characteristics of DR motif antibodies targeting the NA active site through receptor mimicry. **b**, The germline and amino acid sequences of DA03E17 CDR H3 are aligned, with non-templated junction residues in red and residues that have undergone somatic hypermutation in light blue. Nucleotides removed by exonuclease trimming indicated with a line through the letters. Circled bullet points denote antibody main chain contacts only; open circles with rays denote antibody side chain contacts only; Circled bullet points with rays denote both main and side chains contacts. **c**, Search criteria used to identify DR motif precursors from a NGS dataset of 1.1 billion human antibody heavy chain sequences. **d**, Frequency of precursor BCRs with long CDR H3s containing DR (left) or RD (right) motifs, displayed by total and the five D genes with the highest frequencies. Points represent the frequency observed in each of the 14 human donors.

## Discussion

While over 5,000 human mAbs targeting HA have been identified^45^, far fewer antibodies have been characterized for NA^26^. This imbalance underscores the need to continue isolating and studying NA-specific antibodies to better understand the immune response to NA and identify broadly protective epitopes. Such efforts will reveal common molecular features of NA antibody responses that can inform the development of universal vaccines^26^. In this study, we provide a detailed structural and functional analysis of DA03E17, a broadly protective human mAb that targets the NA active site. Using cryo-EM, we determined the structures of DA03E17 bound to NAs from influenza A (H1N1, H3N2) and B viruses, revealing that DA03E17 inhibits the sialidase activity of NA by blocking the highly conserved active site through receptor mimicry using the conserved DR motif in CDR H3. The NA active site is highly conserved, and NA inhibition activity causes arrest of progeny virus egress from the host cell surface preventing spread^60^. Conserved receptor mimicry strategies employed by several NA active site-targeting antibodies likely drive a key mechanism through which broadly protective antibodies inhibit viral spread. Therefore, eliciting antibodies via vaccination that target the NA active site through receptor mimicry holds strong potential for providing broad protection against diverse influenza strains.

DA03E17 retains its binding capacity even in the presence of oseltamivir-resistant mutations, particularly in N1 and N2 subtypes, further demonstrating its robust and broad reactivity. Furthermore, DA03E17 also binds to the NA of the bovine HPAI H5N1 virus currently spreading in U.S. dairy cattle and being transmitted to other mammals, including humans^14^. Considering its strong binding affinity to this bovine H5N1 NA, along with its *in vivo* protective efficacy against the typical avian H5N1 virus (A/Vietnam/1203/2004) isolated from a human, as demonstrated in our previous study^43^, DA03E17 could also serve as a valuable therapeutic option if this virus were to spread more widely in humans.

The ongoing evolution of H3N2 strains and the growing challenge of matching vaccines to circulating viruses^61^ highlight the need for broadly protective antibodies that remain effective against these evolving strains. Our cryo-EM structural analysis and MD simulations revealed that DA03E17 accommodates the N245 glycan of drifted H3N2 virus by inducing conformational changes in the glycan. Similarly, 1G01 induces conformational changes in the glycan itself, while FNI9 requires additional conformational changes in the NA loop to accommodate the N245 glycan. DA03E17 and 1G01 may have an advantage in that they can accommodate this glycan without inducing structural changes in the NA loop, allowing for more stable and efficient binding. The capacity of these antibodies to accommodate the N245 glycan of drifted H3N2 viruses is particularly significant as H3N2-dominated seasons are often more severe, leading to higher hospitalization rates, especially in vulnerable populations^62^.

Our structural and immunogenetic analyses suggest that long CDR H3s with the DR motif near the middle of the CDR H3 could be an important requirement for effectively targeting the NA active site. Bioinformatic analysis using ultradeep NGS data shows that potential precursors with these features are prevalent in the human antibody repertoire. The antibodies compared here have varying angles of approach and peripheral contacts, but they all bind the active site through DR motif-mediated receptor mimicry, maintaining broad reactivities. Additionally, they have diverse germline origins, with IGHJ6 being the only shared feature, indicating their potential to be broadly elicited across human populations. The accompanying manuscript by Lederhofer and Borst et al. further reinforces these findings, presenting additional human antibodies with DR motif-mediated receptor mimicry, thereby underscoring the prevalence of DR motif antibodies in humans, and providing structural and functional insights through both *in vitro* and *in vivo* evaluations. Nevertheless, while the prevalence of potential precursors is promising, the ability of these precursors to mature into functional broadly protective antibodies remains to be fully understood. In conclusion, our study emphasizes the importance of long CDR H3s with the DR motif for targeting the NA active site. The structural insights gained from DA03E17, combined with the investigation of DR motif precursors in the human antibody repertoire, provide a strong foundation for developing vaccines that can elicit broadly protective antibodies. Utilizing this molecular signature to interrogate immune responses to NA and design immunogens capable of triggering protective responses across diverse populations therefore has potential for mitigating future influenza outbreaks.

## Methods

### Expression and purification of recombinant NAs and antibodies

The NAs of the H1N1 A/California/07/2009 (CA09), H1N1 A/Brisbane/02/2018 (BB18), H5N1 A/dairy cattle/Texas/24-008749-001/2024 (TX24), H3N2 A/Perth/16/2009 (PT09), H3N2 A/Indiana/08/2011 (IN11 N2), H3N2 A/Kansas/14/2017 (KS17), H3N2 A/Hong Kong/2671/2019 (HK19), and B/Colorado/06/2017 (B/Victoria-lineage; CO17) were expressed in the mammalian expression system as previously described^46^. Briefly, the ectodomain heads of the IAV N1 (residues 83 to 470 in N2 numbering), N2 (residues 83 to 469 in N2 numbering) NAs, and CO17 B (residues 76 to 466) NA were fused to an N-terminal IL2 signal peptide, a hexahistidine tag, a Strep-tag, a vasodilator-stimulated phosphoprotein (VASP) tetramerization domain, and a thrombin cleavage site. Of note, all N1 NA constructs have previously reported stabilizing mutations in the inter-protomeric interface for the closed tetrameric state (stabilized NA protein, sNAp)^46^. The CA09 N1 sNAp contains ten stabilizing mutations: I99P, Y100L, C161V, E165S, S171A, V176I, S195T, V204I, R419V, and Q412M (N2 numbering). BB18 N1 sNAp and TX24 N1 sNAp also include these ten mutations but carry an additional T131Q substitution. NA constructs were transiently expressed in Expi293F cells (Thermo Fisher Scientific) at 37°C shaking at 125 RPM. The secreted recombinant NA proteins were purified from culture supernatant by Ni-NTA affinity chromatography (Qiagen) followed by size exclusion chromatography using a Superdex 200 Increase 10/300 column (GE Healthcare) in 1X Tris-Buffered Saline (TBS) pH 7.4. The purified NAs were quantified by optical absorbance at 280 nm, and purity and integrity were analyzed by reducing and nonreducing SDS-PAGE. For DA03E17 Fab expression in mammalian cells, the heavy and light chain variable regions of DA03E17 were cloned into the pAbVec containing the corresponding human C_H_1 region of human IgG1 and kappa C_L_ region, respectively. The Fab was transiently expressed in Expi293F cells (Thermo Fisher Scientific) at 37°C shaking at 125 RPM. The secreted recombinant Fab was purified from culture supernatant by CaptureSelect IgG-CH1 Affinity Matrix (Thermo Fisher Scientific) followed by size exclusion chromatography using a Superdex 200 Increase 10/300 column (GE Healthcare) in 1X Tris-Buffered Saline (TBS) pH 7.4.

### ELISA

To evaluate the binding of mAbs, 96-well ELISA plates (PerkinElmer) were coated overnight at 4°C with 2 µg/ml of recombinant NA proteins. Plates were washed three times with a PBS solution containing 0.05% Tween-20 (PBS-T) and then blocked with 5% Non-fat milk (RPI M17200-500.0) in PBS-T. The plates were incubated for 1 hour at room temperature and then washed three times with PBS-T. In the meantime, mAbs were diluted in PBS-T, starting at a concentration of 100 µg/ml with a threefold serial dilution, and then added to the plate for 1 hour at room temperature. After washing, HRP-conjugated goat anti-human IgG Fc secondary antibody (Southern Biotech) was added at a dilution of 1:20,000 in PBS-T and incubated for an additional hour at room temperature. Plates were washed three times with PBS-T and then developed with TMB substrate (Thermo Fisher Scientific). The reaction was stopped by the addition of 2N H_2_SO_4_ and absorbance was measured at 450 nm. The data is shown as one representative biological replicate with the mean ± SD for one ELISA experiment. The ELISAs were repeated 2 times.

### Biolayer Interferometry (BLI)

Antibody affinities were determined by biolayer interferometry using an Octet Red96 (Sartorius). HIS1K biosensors (Sartorius) were hydrated in kinetics buffer (1x PBS, pH 7.4, and 0.002% Tween 20) prior to use. Recombinant NA proteins (10 µg/ml) in kinetics buffer were immobilized on hydrated HIS1K biosensors for 300 sec through their hexahistidine tag at their N-termini and baseline was measured in kinetics buffer for 120 sec. Following baseline measurements, the sensors were loaded with DA03E17 IgG (3-fold serially diluted from 450 nM) and association and dissociation was measured for 300 and 600 sec, respectively. All assay steps were performed at 30 C with agitation set at 1,000 rpm. Baseline correction was carried out by subtracting the measurements recorded for a sensor loaded with the corresponding NA in the same buffer with no Fab. The data were analyzed using the Octet-Red96 software and the association and dissociation rates were calculated using a 1:1 model with global curve fitting.

### Cryo-EM sample preparation and data collection

For each DA03E17 Fab complex with CA09 N1, KS17 N2, and CO17 B NAs, DA03E17 Fab was added at a 1:3 molar ratio of NA protomer to Fab and incubated at room temperature for 1 hour before grid preparation. For the 1G01 IgG-KS17 N2 NA and FNI9 IgG-KS17 N2 NA complexes, each IgG was added at a 1:1 molar ratio of NA tetramer to IgG and incubated at room temperature for 1 hour before grid preparation. Final protein concentrations were between 0.4-0.5 mg/ml. For DA03E17-NA complexes, samples were mixed with octyl-beta-glucoside (OBG; final concentration 0.1% w/v) detergent to aid in particle tumbling and applied to Quantifoil 1.2/1.3 300 mesh copper grids. The grids were plunge-frozen using a Vitrobot Mark IV (Thermo Fisher Scientific) with a blot force of 1 and a blot time of 3.5–5 s at 100% humidity and 4°C. For 1G01- KS17 N2 NA and FNI9-KS17 N2 NA complexes, samples were also mixed with OBG detergent (final concentration 0.1% w/v) but applied to Quantifoil 2/1 400 mesh and gold 1.2/1.3 300 mesh grids. The grids were plunge-frozen at room temperature with 100% humidity, a blot force of 1, and a blot time of 3.5-4.5s. All datasets were collected on a 200 kV Glacios (Thermo Fisher) equipped with a Falcon IV direct electron detector. Automated data collection was carried out using EPU (Thermo Fisher) at a nominal magnification of 190,000× and a pixel size of 0.725 Å, with an approximate exposure dose of 45 e^-^/Å^2^ and a nominal defocus range of -0.7 to -1.4 μm. For each dataset, between 3,116 and 6,416 movie micrographs were collected. For the DA03E17 Fab + KS17 N2 NA dataset, two rounds of data collection resulted in 3,166 and 3,700 movie micrographs, which were combined for processing.

### Cryo-EM data processing

All datasets were processed using cryoSPARC^63^. Dose-weighted movie frame alignment was carried out using Patch motion correction in cryoSPARC live to account for stage drift and beam-induced motion. The contrast transfer function (CTF) was estimated using Patch CTF in cryoSPARC live. Micrographs were curated based on CTF fits, with those worse than 6–8 Å excluded due to poor quality. Individual particles were selected from an initial subset of several hundred micrographs using Blob picker. After several rounds of 2D classification, classes resembling the Fab-NA complex were used to template pick for all micrographs. Clean particle stacks were selected through multiple rounds of 2D classification and used to generate a reference volume via ab initio reconstruction, which was then utilized as an initial volume for homogeneous refinement and/or non-uniform refinement. For the DA03E17-CA09 N1 sNAp and DA03E17-KS17 N2 NA datasets, Fab-NA complex particles underwent 3D classification. The best 3D classes were refined using either local refinement with a Fab-NA mask (DA03E17-CA09 N1) or non-uniform refinement (DA03E17-KS17 N2). Final maps were generated after global CTF refinement and application of C4 symmetry. For the apo-KS17 N2 NA, only NA particles without visible Fab binding were selected through multiple rounds of 2D classification, followed by non-uniform refinement with C4 symmetry applied and global CTF refinement, yielding the final map. For the DA03E17-CO17 B NA dataset, Fab-NA complex particles were subjected to homogeneous refinement with C4 symmetry applied, followed by global CTF refinement to generate the final map. For the 1G01-KS17 N2 NA dataset, particles selected after 2D classification were subjected to heterogeneous refinement. The best 3D classes were refined using local refinement with a Fab-NA mask and C4 symmetry applied, yielding the final map. For the FNI9-KS17 N2 NA dataset, particles selected after 2D classification were subjected to ab initio reconstruction with five classes. Particles from the highest-quality classes were then used for homogeneous refinement and local refinement with a Fab-NA mask, with no symmetry applied, yielding the final map.

### Cryo-EM model building and refinement

For model building, the structures of CA09 N1 NA (PDB: 6Q23), HK19 N2 NA (PDB: 8G3O), and PK13 B NA (PDB: 6V4N) were used as the initial models for CA09 N1 sNAp, KS17 N2 NA, and CO17 B NA, respectively. The DA03E17 Fv model was generated using ABodyBuilder2^64^. The models were fitted into the cryo-EM maps using UCSF ChimeraX^65^. The models were manually adjusted using Coot^66^ and further refined through Rosetta Relax^67^ and real-space refinement in Phenix^68^. N1 and N2 NAs were numbered according to the N2 numbering scheme, while the antibody Fv was numbered based on the Kabat numbering scheme. Epitope and paratope residues, as well as their interactions, were identified by using the PISA server^69^ with buried surface area (BSA >5 Å^2^) as the criterion. Structural figures were prepared using UCSF ChimeraX.

### NA Sequence conservation analysis

For Extended Data Table 1, NA sequences for IAV Group 1 (N1, N4, N5, and N8), Group 2 (N2, N3, N6, N7, and N9), and IBV (Victoria and Yamagata) were retrieved from GISAID (www.gisaid.org). Sequences for each subtype were aligned using GENETYX ver13, and SNP analysis was conducted using the BV-BRC platform (https://www.bv-brc.org/). For Figure 2, full-length human IAV H1N1 and H3N2 and IBV Victoria-lineage NA protein sequences circulating from 1977 to 2023, 1968 to 2023, and 1987 to 2023, respectively, were downloaded from GISAID. To avoid temporal sampling bias, we sampled at most 10 sequences per year, which resulted in total 340, 421, and 278 sequences for N1, N2, and B/Victoria NAs. The multiple sequence alignments were analyzed with Clustal Omega and the respective references used for alignment were H1N1 A/California/04/2009, H3N2 A/Kansas/14/2017 and B/Colorado/06/2017 for N1, N2 and B/Victoria NAs, respectively. NA sequences used in the analyses were listed in Item S1. Sequence logos were generated by WebLogo 3 and manually curated in Adobe Illustrator.

### Molecular Dynamics

As starting models for our MD simulations, the cryo-EM structures of DA03E17-KS17 N2 NA (this study) and apo-KS17 N2 NA (this study) were used. The starting structures for our simulations were prepared in Molecular Operating Environment (MOE, CCG)^70^ using the Protonate3D tool^71^. To neutralize the charges, the uniform background charge was applied, which is required to compute long-range electrostatic interactions^72^. Using the tleap tool of the AmberTools22^73^ package, the structures were soaked in cubic water boxes of TIP3P water molecules with a minimum wall distance of 12 Å to the protein^74–76^. For all simulations, parameters of the AMBER force field 19SB were used^77^. For the glycans we used the most recent GLYCAM force field, namely the GLYCAM-06j parameter set^78^. We then performed 3 repetitions of 1 µs of classical molecular dynamics simulations for each system. Molecular dynamics simulations were performed in an NpT ensemble using pmemd.cuda^79^. Bonds involving hydrogen atoms were restrained by applying the SHAKE algorithm^80^ allowing a time step of 2 fs. The Langevin thermostat was used to maintain the temperature during simulations at 300 K^81,82^ with a collision frequency of 2 ps^−1^ and a Monte Carlo barostat^83^ with one volume change attempt per 100 steps. Clustering analysis has been performed using the hierarchical average linkage clustering implement in cpptraj, by aligning on the Cα-positions of the NA and clustering on the glycans using a RMSD distance cut-off criterion of 2.5 Å^84^.

### Antibody precursor frequency estimates

Precursor searching and frequency estimates of NA active site-targeting antibodies with receptor mimicry were performed using methods described previously (Ma et al., 2024, under review). The precursor definition had two requirements: (1) CDR H3 length ranging from 19 to 30 amino acids, and (2) the DR or RD motif occurred approximately in the middle of the CDR H3, ≥ 8 amino acids on either side. These criteria were based on the CDR H3 length and DR/RD motif position in several NA active site-targeting antibodies. These definitions allowed for precursors with long CDR H3s possessing a DR or RD motif, regardless of germline gene usage, with diverse V-D and D-J junctions. A total of 1.1 x 10^9^ BCR heavy chain sequences were searched in a ultradeep NGS dataset of 14 human donors (PMID: 30664748 and 31672916). PySpark scripts used in this analysis are available at https://github.com/SchiefLab/Jo2024 along with instructions on setting up an EMR cluster.

## Data availability

The atomic models and cryo-EM density maps generated in this study have been deposited to the Protein Data Bank (PDB) and Electron Microscopy Databank (EMDB) respectively. The accession codes are 9CYE and EMD-46041 (DA03E17 Fab+CA09 N1 NA), 9CYF and EMD- 46042 (DA03E17 Fab+KS17 N2 NA), 9CYG and EMD-46043 (Apo-KS17 N2 NA), 9CYH and EMD-46044 (DA03E17 Fab+CO17 B NA), 9CYI and EMD-46045 (1G01 Fab+KS17 N2 NA), and 9CYJ and EMD-46046 (FNI9 Fab+KS17 N2 NA).

## Acknowledgments

We thank Hannah Turner and Will Lessin for electron microscopy support. We thank Charles Bowman and JC Ducom for computational support. We thank Lauren Holden for administrative support. This research was supported by the NIH National Institute of Allergy and Infectious Diseases (NIAID) Collaborative Influenza Vaccine Innovation Centers (CIVICs) contract grant 75N93019C00051, Third Rock Ventures, Basic Science Research Program through the National Research Foundation of Korea (NRF) funded by the Ministry of Education (RS-2023-00240483), and the Japan Agency for Medical Research and Development (JP24wm0125002 and JP243fa627001).

## Author Contributions

J. Han and A.B.W. conceptualized the studies. G.J., J. Huang, J.C., and A.J.R. expressed and purified protein. G.J., J.L.T., J.A.F., and J. Han prepared cryo-EM samples, and collected and processed cryo-EM data. G.J. built and refined atomic models. G.J. performed biochemical experiments. G.J., S.Y., and O.S. conducted the NA sequence conservation analysis. J.M.S. and K.M.M. performed the precursor frequency analysis. M.L.F.Q. performed molecular dynamics simulations. G.J., J.M.S., K.M.M., S.Y., and M.L.F.Q. wrote the original manuscript draft with input and edits from Y.K., J. Han, and A.B.W. All authors contributed to the manuscript review and editing.

