## Supplemental Material for "Structural basis of broad protection against influenza virus by a human antibody targeting the neuraminidase active site via a recurring motif in CDR H3"

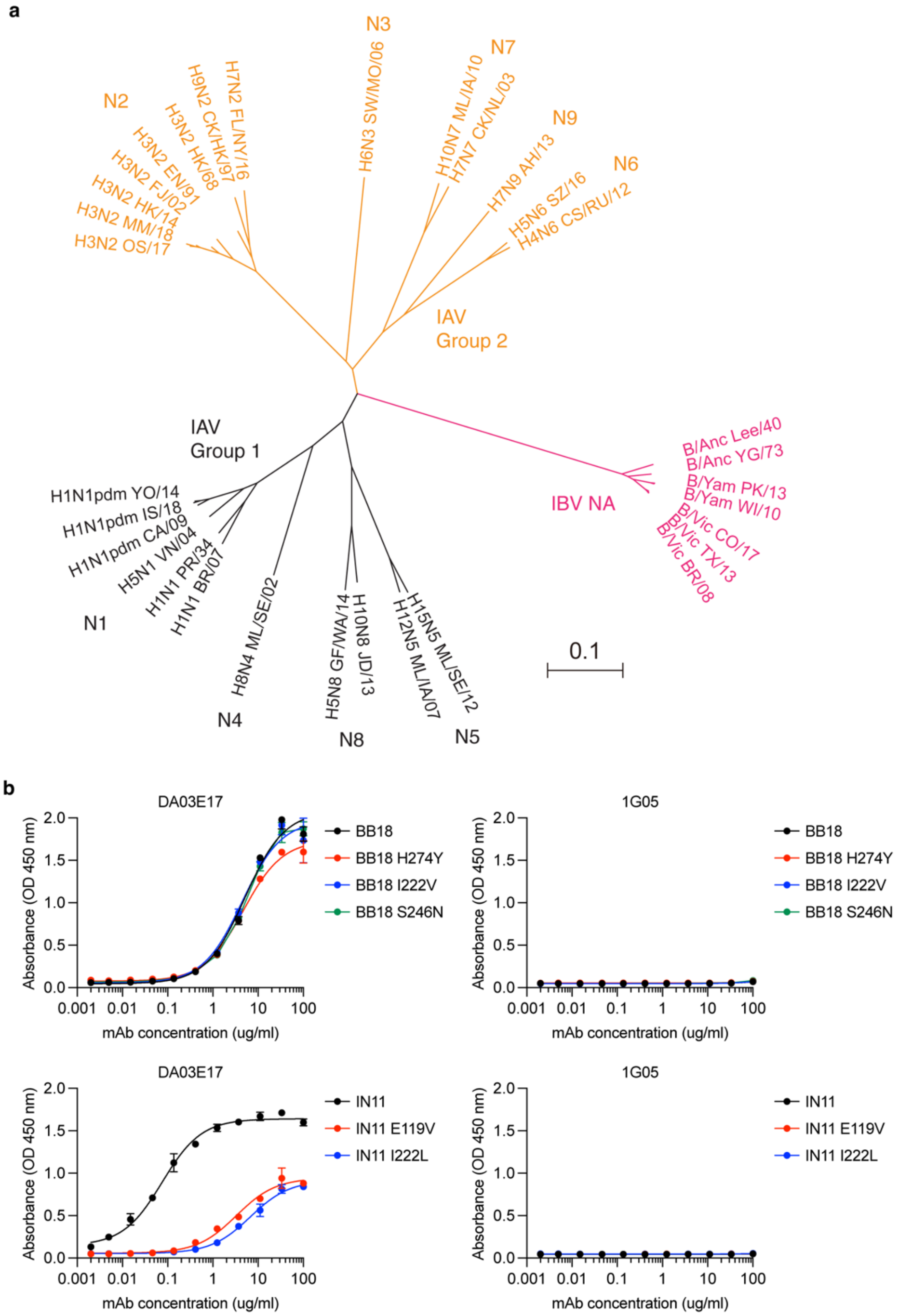


**Extended Data Fig.1 Breadth of mAb DA03E17. a**, Phylogenetic tree of influenza A and B virus NAs that DA03E17 showed reactivities (adapted from our previous study[1]). The scale bar represents a 10% change in amino acids. Complete viral strain designations are listed in Source Data file. **b**, ELISA binding curves of DA03E17 against recombinant NAs from H1N1 A/Brisbane/02/2018 (BB18 N1 sNAp) and H3N2 A/Indiana/08/2011 (IN11 N2) with or without the oseltamivir-resistant mutations. Influenza B virus NA-specific mAb 1G05 was used as negative control. Data are plotted as mean ± SD.

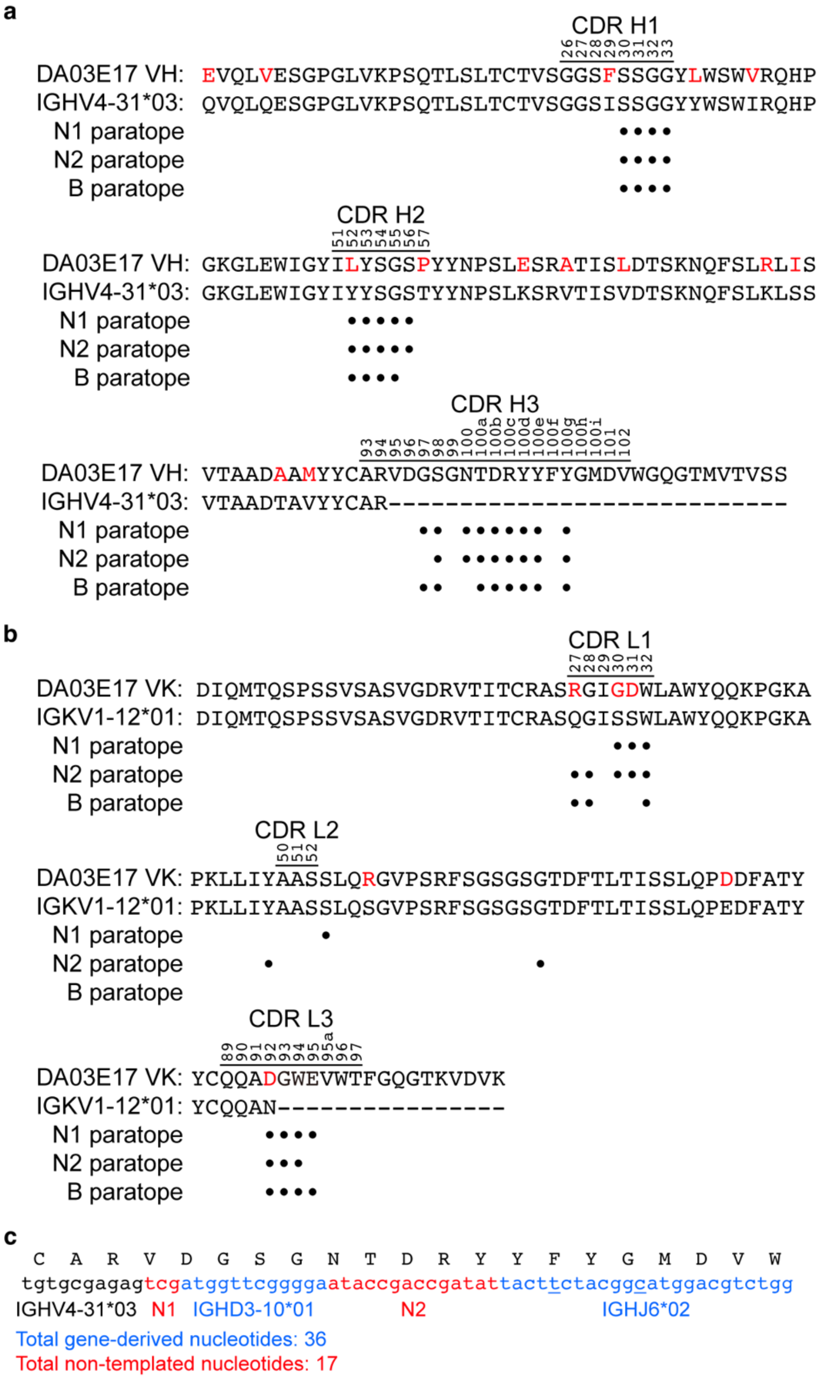


**Extended Data Fig.2 Comparison of DA03E17 sequence to its putative germline sequence.** **a**,**b**, Alignment of the DA03E17 variable region sequence with the germline IGHV4-31*03 sequence (**a**), and with the germline IGKV1-12*01 sequence (**b**). The regions that correspond to CDRs are indicated based on the IMGT CDR definition scheme. Residues that differ from the germline are highlighted in red. Residues that interact with each NA are indicated with black dot. **c**, Sequence of VDJ junction of DA03E17 is shown with putative germline gene segments as well as non-templated N-regions indicated. Somatically mutated nucleotides are underlined.

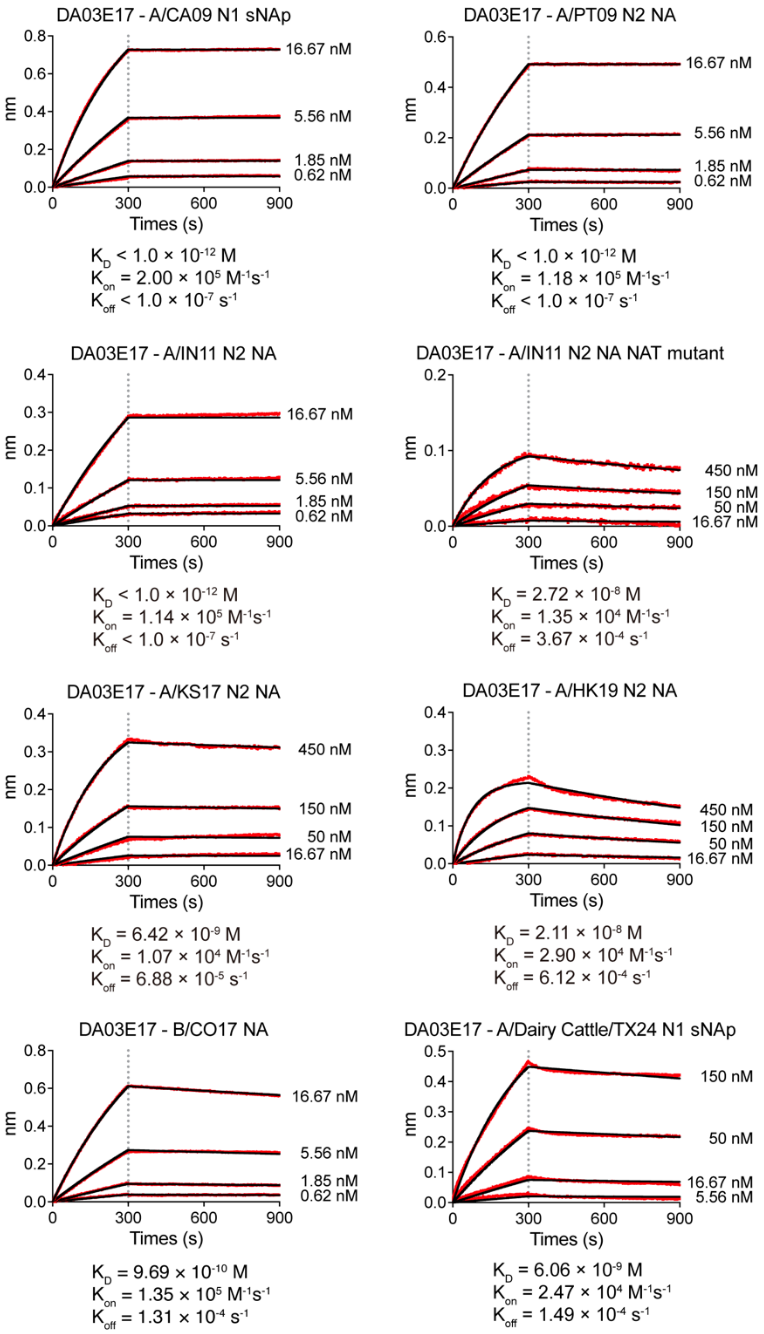


**Extended Data Fig.3 Binding kinetics of DA03E17.** Binding of the DA03E17 IgG to recombinant NAs from H1N1 A/California/07/2009 (A/CA09 N1 sNAp), H3N2 A/Perth/16/2009 (A/PT09 N2 NA), H3N2 A/Indiana/08/2011 (A/IN11 N2 NA), H3N2 A/Indiana/08/2011 S245N/S247T mutant (A/IN11 N2 NA NAT mutant), H3N2 A/Kansas/14/2017 (A/KS17 N2 NA), H3N2 A/Hong Kong/2671/2019 (A/HK19 N2 NA), B/Colorado/06/2017 (B/Victoria-lineage; B/CO17 NA), and HPAI H5N1 clade 2.3.4.4b A/Dairy cattle/Texas/24-008749-001/2024 (A/Dairy cattle/TX24 N1 sNAp) as determined by bio-layer interferometry. The experimental data are shown in red, and the fitted data are shown in black. For estimating the $K_D$, a 1:1 binding model was used.

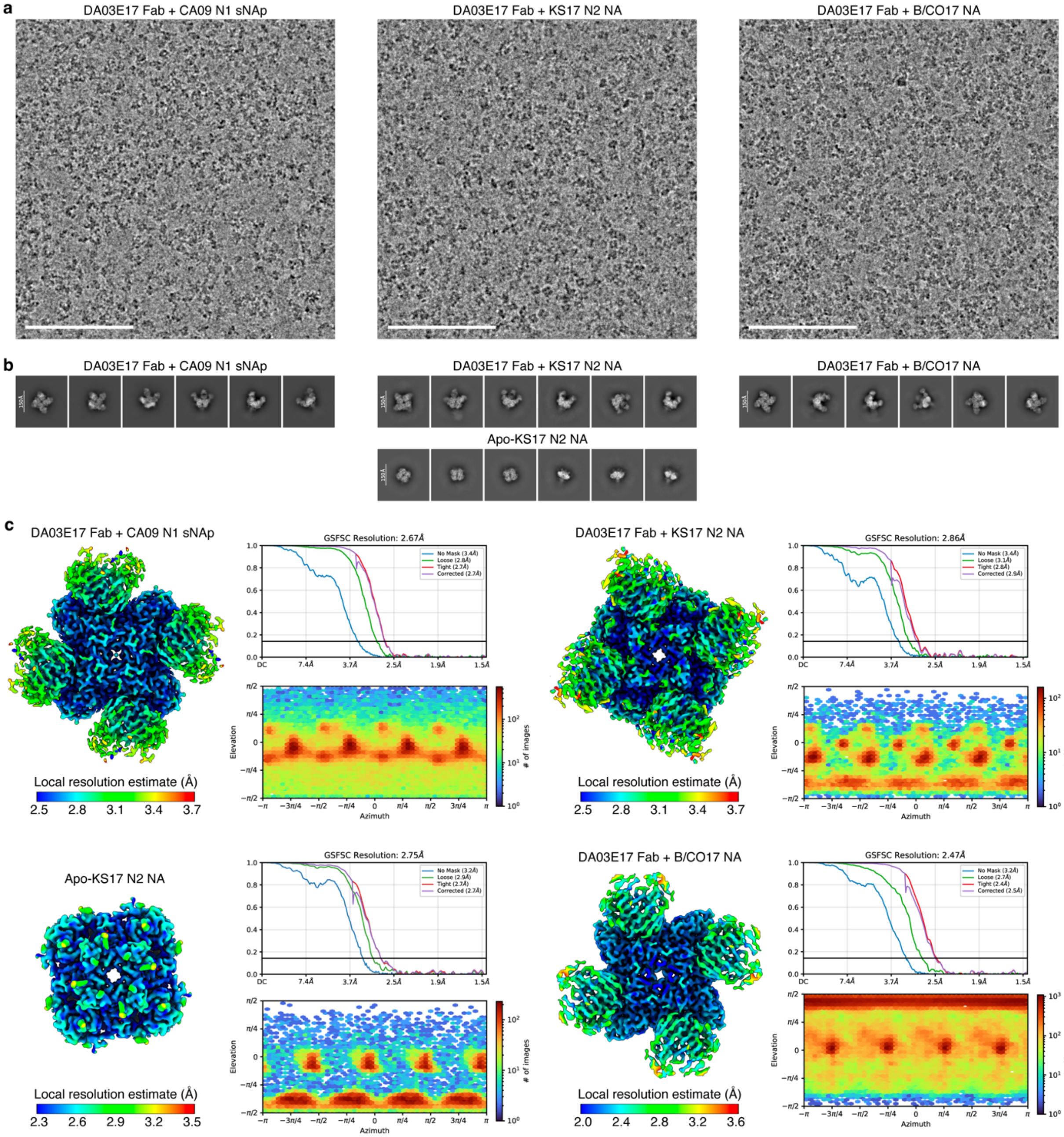


**Extended Data Fig.4 Cryo-EM data processing and validation of DA03E17 in complex with CA09 N1 sNAp, and KS17 N2 and CO17 B NAs. a**, Representative micrographs of DA03E17 Fab in complex with CA09 N1 sNAp, and KS17 N2 and CO17 B NAs. Scale bar, 100 nm. **b**, Representative 2D class averages. Scale bar, 150 Å. **c**, Local resolution maps, gold-standard Fourier shell correlation curves, and viewing direction distributions. The 0.143 cutoff is indicated by a horizontal black line.

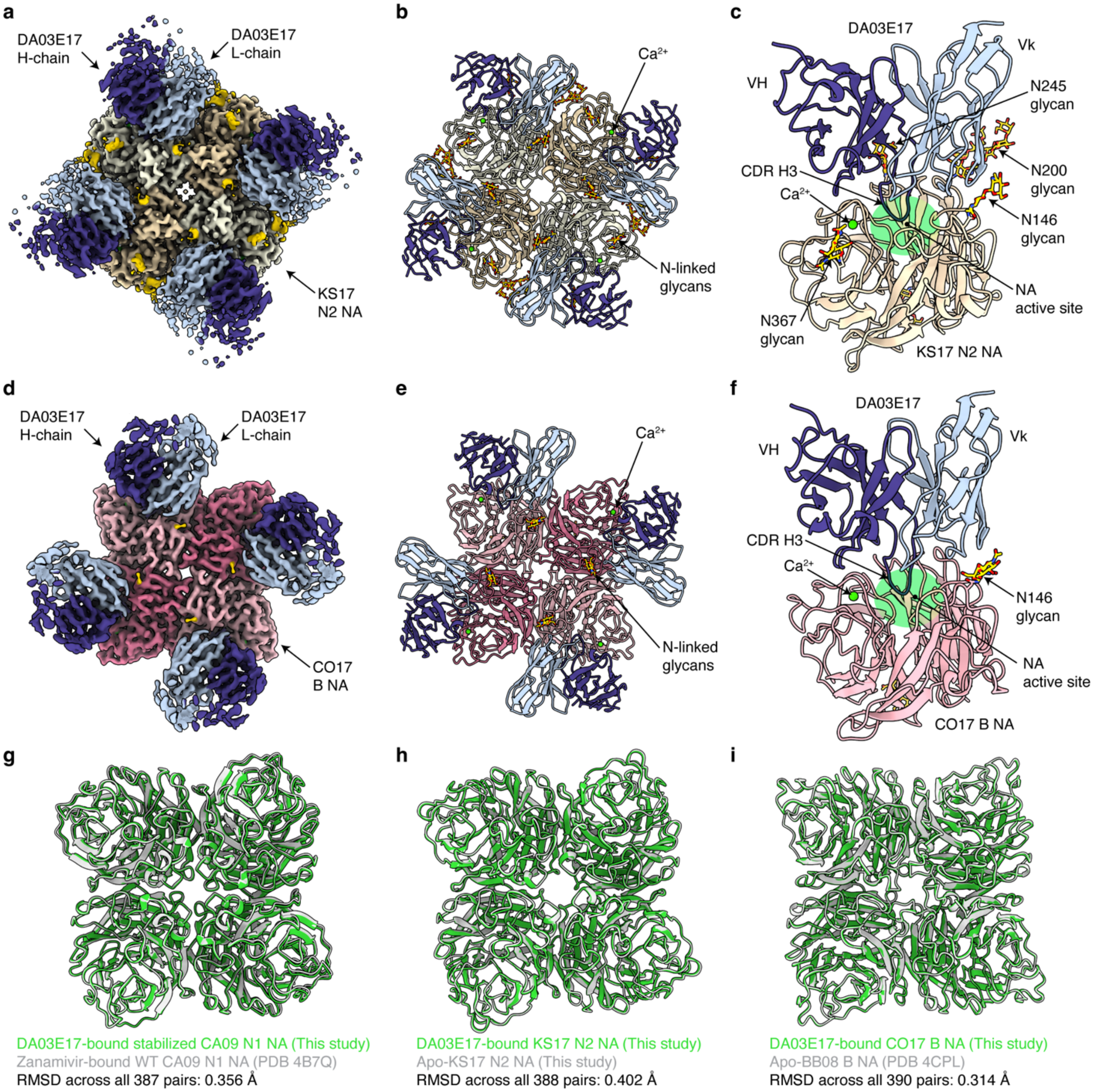


**Extended Data Fig.5 Cryo-EM structures of DA03E17 Fab in complex with KS17 N2 and CO17 B NAs. a**,**b**, Overall structure of DA03E17 Fab and KS17 N2 NA complex. Cryo-EM map at 2.86 Å (**a**) and atomic model (**b**) from top view. **c**, Ribbon diagram of the KS17 N2 NA protomer bound with one DA03E17 Fab. **d**,**e**, Overall structure of DA03E17 Fab and CO17 B NA complex. Cryo-EM map at 2.47 Å (**d**) and atomic model (**e**) from top view. **f**, Ribbon diagram of the CO17 B NA protomer bound with one DA03E17 Fab. **g**–**i**, Structural comparison between DA03E17-bound N1 sNAp (**g**) and N2 (**h**) and B (**i**) NAs and corresponding WT NAs.

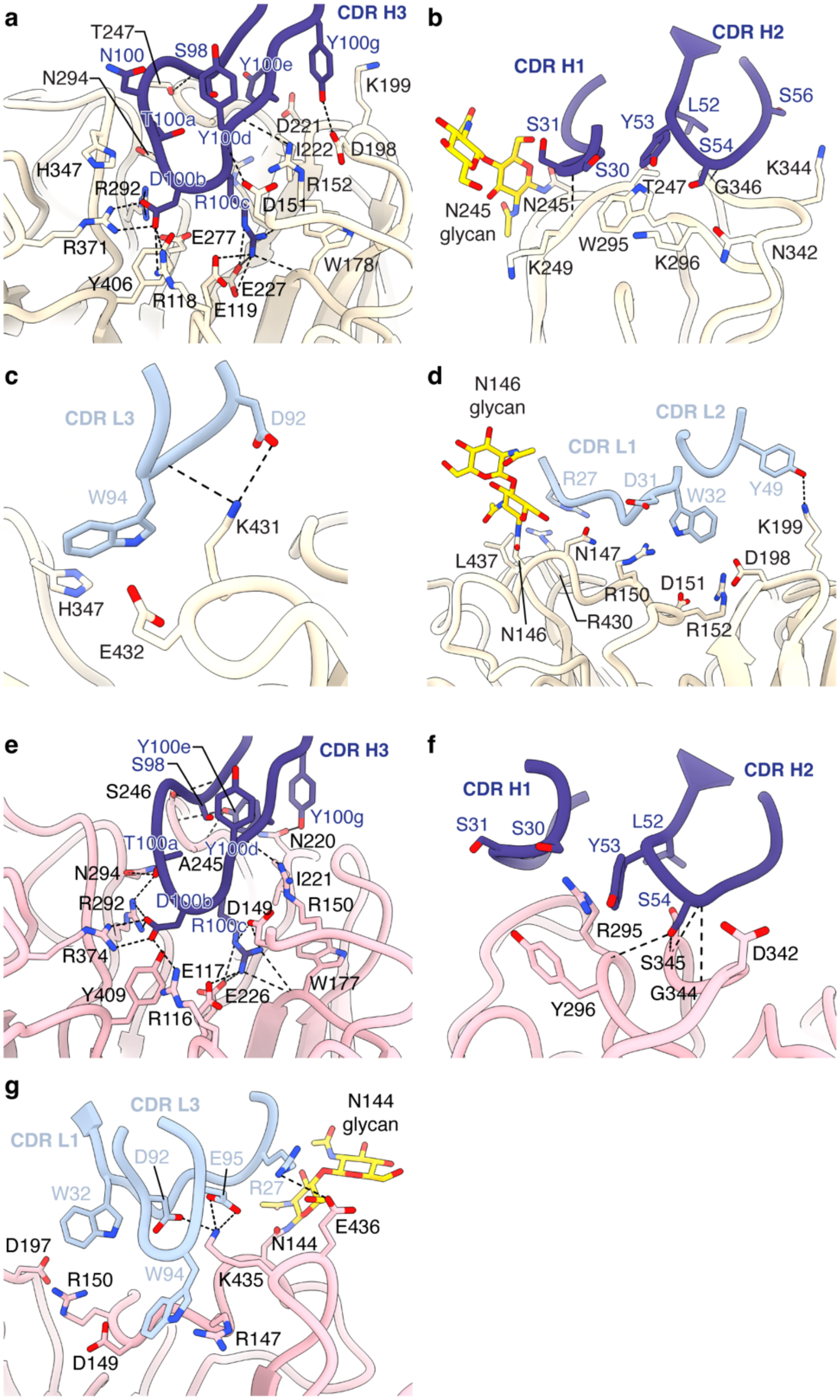


**Extended Data Fig.6 Interactions between DA03E17 and KS17 N2 and CO17 B NAs. a–d**, Detailed illustrations of the interactions between KS17 N2 NA and DA03E17 CDR H3 (**a**), CDRs H1 and H2 (**b**), CDR L3 (**c**), and CDRs L1 and L2 (**d**). **e–g**, Detailed illustrations of the interactions between CO17 B NA and DA03E17 CDR H3 (**e**), CDRs H1 and H2 (**f**), and CDRs L1 and L3 (**g**).

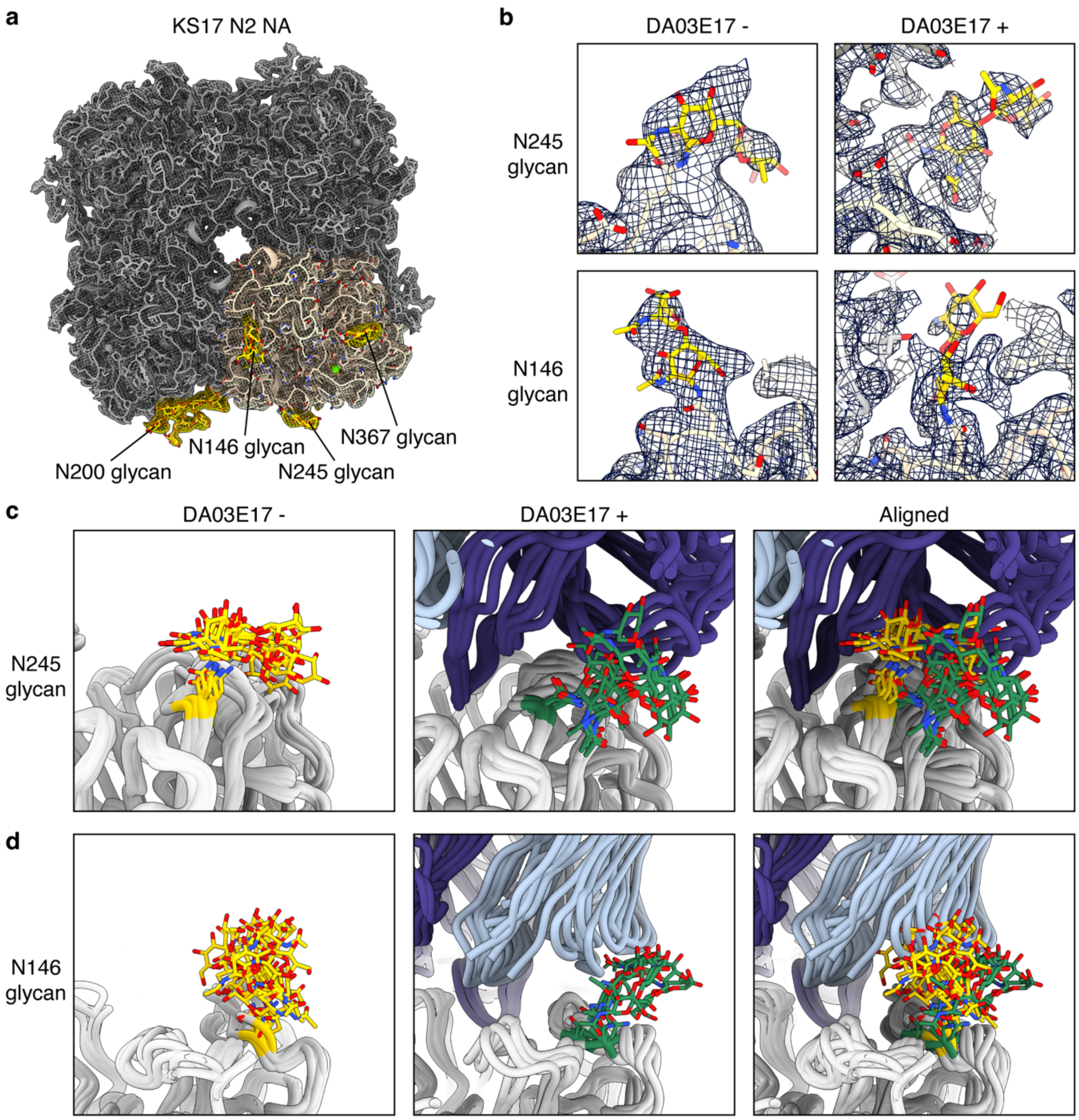


**Extended Data Fig.7 Molecular dynamics simulations of DA03E17-KS17 N2 NA complex. a**, Structure of KS17 N2 NA in apo-form with overlaid electron density map. The glycans of one protomer are highlighted in yellow. **b**, Electron density maps (blue mesh) and modeled glycan structures of the N245 and N146 glycans (yellow) in the app-KS17 N2 NA (left) and DA03E17-KS17 N2 NA complex (right) structures, showing conformational changes upon DA03E17 binding. **c**–**d**, Cluster representatives of KS17 N2 NA from MD simulations, shown without DA03E17 bound and with DA03E17 bound, highlighting the structural diversity of the N245 (**c**) and N146 (**d**) glycans in the free state and the population shift of these glycans upon DA03E17 binding.

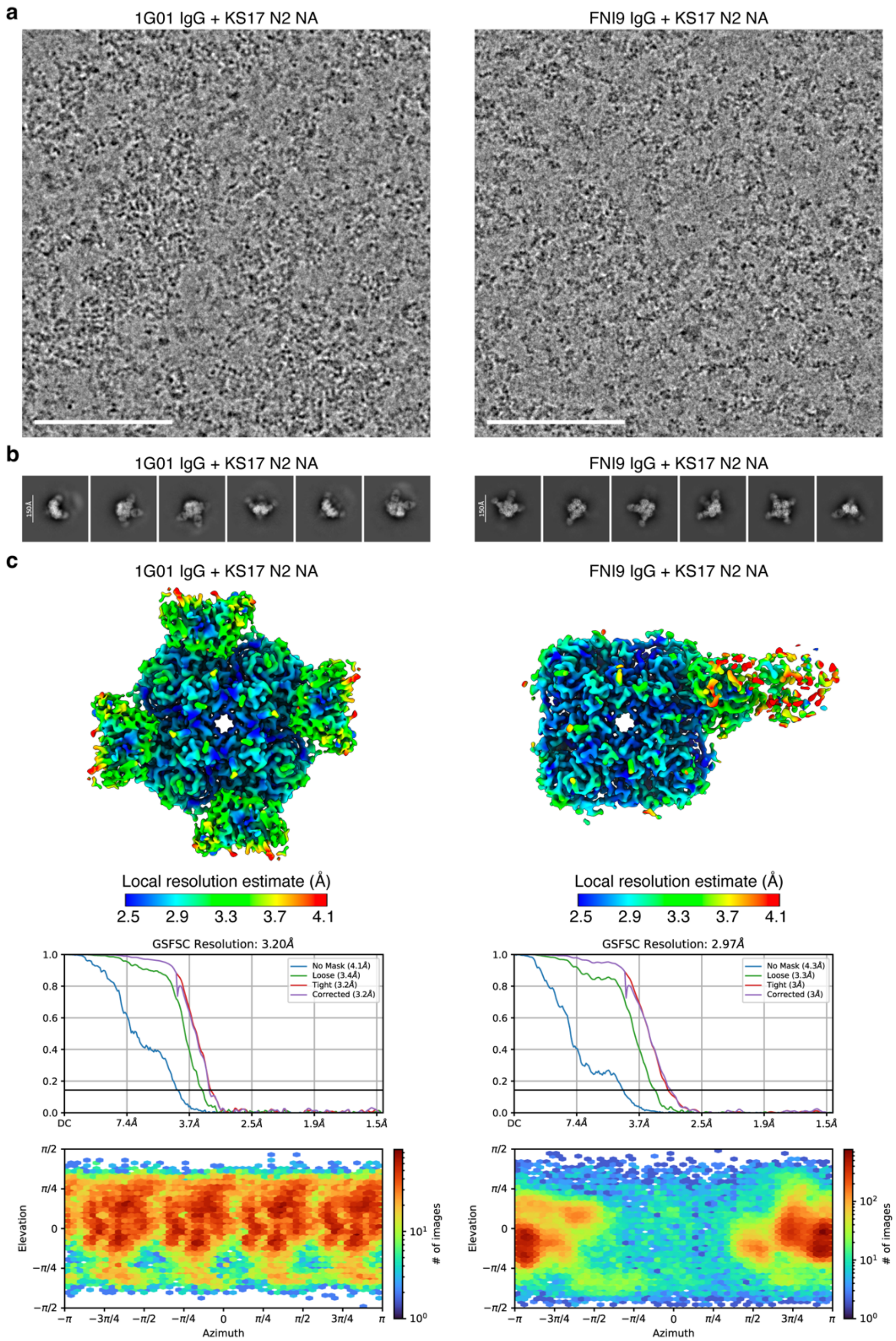


**Extended Data Fig.8 Cryo-EM data processing and validation of KS17 N2 NA in complex with previously reported broadly protective antibodies, 1G01 and FNI9. a**, Representative micrographs of KS17 N2 NA in complex with 1G01 IgG and FNI9 IgG. Scale bar, 100 nm. **b**, Representative 2D class averages. Scale bar, 150 Å. **c**, Local resolution maps, gold-standard Fourier shell correlation curves, and viewing direction distributions. The 0.143 cutoff is indicated by a horizontal black line.

### Extended Data Table 1 DA03E17 epitope conservation across IAV subtypes and IBV lineages.

| Met=1 numbering | | | DA03E17 CA09 N1 epitope | DA03E17 KS17 N2 epitope | DA03E17 CO17 B epitope | IAV Group 1 | | | | IAV Group 2 | | | | | IBV | |
|---|---|---|---|---|---|---|---|---|---|---|---|---|---|---|---|---|
| N1 | N2 | B | | | | N1 (98,463)[a] | N4 (484) | N5 (752) | N8 (6,667) | N2 (135,160) | N3 (2,187) | N6 (5,727) | N7 (1,558) | N9 (3,447) | Victoria lineage (30,156) | Yamagata lineage (14,098) |
| 118 | 118 | 116 | R | R | R | R[b] (99.99)[c] | R (99.59) | R (99.87) | R (99.96) | R (99.99) | R (100) | R (99.98) | R (100) | R (99.97) | R (99.99) | R (99.99) |
| 119 | 119 | 117 | E | E | E | E (99.67) | E (100) | E (99.87) | E (99.97) | E (99.92) | E (99.95) | E (99.74) | E (100) | E (99.59) | E (99.98) | E (99.96) |
| 146 | 146 | 144 | - | N | N | N (99.80) | N (100) | N (100) | S (99.60) | N (95.34) | N (99.68) | N (99.97) | N (100) | N (99.97) | N (99.92) | N (99.93) |
| 147 | 147 | 145 | - | N | - | G (99.81) | G (100) | N (99.87) | N (99.93) | N (82.72) | G (99.91) | G (99.32) | G (99.42) | G (99.71) | G (99.88) | G (99.91) |
| 149 | 149 | 147 | I | - | R | I (77.68) | V (99.59) | V (99.34) | V (96.24) | V (82.57) | I (85.56) | I (96.68) | I (97.43) | I (98.99) | R (99.95) | R (99.94) |
| 150 | 150 | 148 | K | R | - | K (99.70) | K (99.79) | K (99.87) | K (99.39) | R (77.34) | K (98.58) | H (97.59) | H (99.42) | H (99.62) | G (98.93) | E (96.71) |
| 151 | 151 | 149 | D | D | D | D (99.01) | D (100) | D (100) | D (99.97) | D (95.78) | D (99.31) | D (99.97) | D (100) | D (99.94) | D (99.64) | D (99.77) |
| 152 | 152 | 150 | R | R | R | R (99.96) | R (100) | R (100) | R (99.91) | R (99.98) | R (99.95) | R (99.83) | R (99.94) | R (99.86) | R (100) | R (99.98) |
| 179 | 178 | 177 | W | W | W | W (99.99) | W (100) | W (100) | W (99.93) | W (99.88) | W (100) | W (99.98) | W (100) | W (100) | W (100) | W (99.99) |
| 199 | 198 | 197 | D | D | D | D (99.74) | D (100) | D (100) | D (99.96) | D (99.91) | D (100) | N (96.26) | N (100) | N (99.59) | D (99.82) | D (99.51) |
| 200 | 199 | 198 | N | K | - | S (53.88) | S (51.34) | D (96.54) | S (56.81) | K (89.04) | N (97.58) | N (96.89) | D (93.77) | N (99.97) | N (99.40) | S (95.39) |
| 222 | 221 | 220 | N | D | N | N (67.75) | N (98.76) | Q (99.47) | D (99.31) | D (71.66) | D (86.69) | N (99.56) | N (99.61) | N (99.19) | N (93.03) | N (97.43) |
| 223 | 222 | 221 | I | I | I | I (99.68) | I (98.56) | I (100) | I (99.81) | I (99.65) | I (98.95) | I (99.91) | I (99.94) | I (99.71) | I (99.92) | I (99.82) |
| 225 | 224 | 223 | R | R | R | R (99.98) | R (100) | R (99.87) | R (99.97) | R (99.97) | R (99.82) | R (99.97) | R (100) | R (100) | R (99.99) | R (99.94) |
| 228 | 227 | 226 | E | E | E | E (99.97) | E (100) | E (99.87) | E (99.99) | E (99.98) | E (100) | E (99.97) | E (99.94) | E (100) | E (99.99) | E (99.99) |
| 246 | 245 | 244 | P | N | S | P (99.77) | P (100) | P (100) | P (99.96) | N (65.54) | P (99.95) | P (98.67) | S (98.59) | S (62.98) | S (99.72) | P (95.21) |
| 247 | 246 | 245 | S | A | A | S (99.52) | S (99.79) | A (99.87) | A (81.97) | A (99.88) | A (99.95) | A (99.83) | A (100) | A (99.74) | A (99.93) | A (99.93) |
| 248 | 247 | 246 | N | T | S | D (74.86) | D (98.14) | N (89.64) | N (99.57) | T (66.24) | A (86.47) | N (97.96) | S (97.11) | T (99.83) | S (99.83) | S (98.25) |
| 249 | 248 | 247 | G | G | G | G (97.46) | A (99.79) | N (72.91) | R (98.64) | G (99.55) | N (86.15) | N (89.70) | S (97.88) | G (100) | G (99.63) | G (99.87) |
| 250 | 249 | 248 | - | K | - | Q (96.02) | Q (100) | Q (99.87) | Q (99.91) | K (91.12) | S (84.92) | R (57.48) | Q (99.74) | P (100) | V (98.74) | I (88.11) |
| 277 | 276 | 275 | - | - | E | E (99.99) | E (100) | E (99.87) | E (99.99) | E (99.92) | E (100) | E (99.98) | E (100) | E (100) | E (99.97) | E (99.92) |
| 278 | 277 | 276 | E | E | E | E (99.99) | E (100) | E (99.87) | E (99.99) | E (99.99) | E (100) | E (99.91) | E (100) | E (100) | E (99.98) | E (99.91) |
| 293 | 292 | 292 | R | R | R | R (99.98) | R (100) | R (100) | R (99.96) | R (99.96) | R (99.95) | R (99.97) | R (100) | R (98.17) | R (99.98) | R (99.95) |
| 295 | 294 | 294 | N | N | N | N (99.93) | N (100) | N (99.73) | N (99.97) | N (99.98) | N (99.95) | N (99.93) | N (99.94) | N (99.97) | N (99.98) | N (99.94) |
| 296 | 295 | 295 | W | W | R | W (99.99) | W (100) | W (100) | W (100) | W (99.98) | W (100) | W (99.97) | W (100) | W (100) | R (92.77) | S (85.10) |
| 297 | 296 | 296 | H | K | Y | H (99.98) | R (97.32) | N (99.60) | T (80.26) | K (99.00) | K (99.73) | K (97.44) | Q (99.94) | Q (99.36) | Y (99.96) | Y (99.88) |
| 339 | 342 | 341 | S | N | D | S (85.99) | N (98.56) | G (100) | G (99.99) | N (99.98) | S (99.41) | T (92.10) | T (99.42) | P (99.97) | D (96.15) | D (94.54) |
| 341 | 344 | 343 | N | K | K | N (95.20) | K (99.79) | T (100) | Deletion | K (48.55) | N (86.71) | S (85.81) | P (99.94) | N (97.65) | K (59.91) | E (89.02) |
| 342 | 345 | 344 | G | G | G | G (99.99) | E (100) | N (98.54) | Q (89.79) | G (99.98) | G (99.95) | P (97.64) | G (99.36) | N (97.80) | G (99.97) | G (99.95) |
| 343 | 346 | 345 | A | G | S | A (99.54) | R (99.79) | N (99.87) | G (99.30) | G (71.59) | G (85.39) | D (98.12) | A (97.11) | N (99.97) | S (98.78) | S (98.94) |
| 344 | 347 | 346 | N | H | - | N (85.31) | Y (100) | Y (99.73) | Y (99.90) | H (92.20) | P (85.30) | P (99.88) | P (99.74) | N (98.98) | G (98.74) | G (99.41) |
| 368 | 371 | 374 | R | R | R | R (99.98) | R (100) | R (99.87) | R (99.66) | R (99.99) | R (100) | R (99.97) | R (100) | R (99.94) | R (100) | R (99.98) |
| 402 | 406 | 409 | Y | Y | Y | Y (99.98) | Y (100) | Y (99.87) | Y (99.63) | Y (99.99) | Y (100) | Y (99.97) | Y (100) | Y (99.97) | Y (100) | Y (99.99) |
| 430 | 430 | 433 | R | R | - | R (91.58) | Q (100) | K (99.47) | K (52.17) | R (99.49) | R (84.11) | R (98.62) | R (99.36) | R (92.95) | G (99.80) | G (99.89) |
| 431 | 431 | 434 | P | K | - | P (99.89) | P (99.79) | P (99.47) | P (100) | K (90.95) | P (99.68) | P (99.84) | P (100) | P (99.97) | G (99.93) | G (99.95) |
| 432 | 432 | 435 | K | E | K | E (61.35) | K (99.79) | E (98.94) | E (99.85) | E (87.68) | N (99.77) | K (96.28) | E (100) | K (99.51) | K (99.79) | K (99.14) |
| 433 | 433 | 436 | - | - | E | E (99.90) | E (100) | E (100) | E (99.97) | E (99.92) | K (89.22) | E (99.83) | E (100) | E (99.94) | E (99.49) | T (95.42) |
| 436 | 437 | 437 | - | L | - | I (97.77) | I (99.79) | I (100) | I (99.60) | L (83.37) | S (99.27) | L (97.84) | W (98.78) | W (99.80) | T (99.93) | T (99.83) |

DA03E17 epitope residues conserved across influenza A and B viruses are highlighted in cyan. Epitope residues highly conserved (>90%) within the corresponding IAV group or IBV and within the corresponding subtype or lineage are highlighted in green and yellow, respectively.

[a] Number of influenza NA sequences for each subtype or lineage obtained from the GISAID database (https://www.gisaid.org/) in November 2023.

[b] Most common residue at each position.

[c] Percent identity for most common residue

1. Yasuhara, A. et al. A broadly protective human monoclonal antibody targeting the sialidase activity of influenza A and B virus neuraminidases. *Nat Commun* **13**, 6602 (2022).
